# Loss of Mitochondrial Respiratory Capacity Reshapes Myeloid Cell Function during *Mycobacterium tuberculosis* infection

**DOI:** 10.64898/2026.09.04.749543

**Authors:** Eduardo L. Martinez, Cory J. Mabry, Aja K. Coleman, Morgan S. Osborne, Joshua Simmons, Stacy Hahn, Jessica B. Huskey, Kaitlyn S. Armijo, Taylor Newbolt, Jacob R. Davis, Lianna Zhou, Spyros Kalams, Denis Mogilenko, Kristin L. Patrick, Robert O. Watson

## Abstract

Myeloid cells are essential mediators of host defense against *Mycobacterium tuberculosis* (Mtb), yet the metabolic programs that sustain their function during chronic infection remain poorly defined. Here, using scRNA-seq we identified a striking, coordinated decline in mitochondrial electron transport chain gene expression across diverse myeloid populations as Mtb disease progressed in mice. This transcriptional remodeling was associated with broad changes in immune and metabolic pathways, including reduced antigen presentation, interferon responses, protein synthesis, and glycolysis. Accordingly, loss of Complex I in macrophages (*Ndufs4* knockdown) reduced MHC-II surface expression, dysregulated inflammatory gene expression, and limited control of Mtb replication. Finally, analysis of single-cell transcriptomic data from Mtb-exposed human household contacts identified an almost identical transcriptional program enriched in IGRA+ individuals, supporting a role for mitochondrial respiratory remodeling in human TB. Together, these findings demonstrate that mitochondrial bioenergetic competence is required to sustain macrophage effector function during chronic Mtb infection and suggest that mitochondrial restoration may boost protective responses in TB patients.

## INTRODUCTION

During *Mycobacterium tuberculosis* (Mtb) infection, myeloid cells function both as critical mediators of antimicrobial immunity and cellular niches for bacterial persistence (Warner et al., 2025). Macrophages, monocytes, and dendritic cells contribute to bacterial restriction through inflammatory signaling, antimicrobial activity, antigen processing and presentation, and coordination of adaptive immunity. Because Mtb can establish a long term-persistent infection, myeloid cells must function within a chronically inflamed, hypoxic, and nutrient-limited environment within the tissue environment (Babunovic et al., 2022; Cadena et al., 2017; Liu et al., 2025; Samstein et al., 2013; Srivastava & Ernst, 2014). Effective control of Mtb therefore requires sustained metabolic adaptation that preserves antimicrobial function, including cytokine production, phagolysosomal activity, antigen presentation, and T-cell support. However, this response must also remain appropriately regulated: although a weak immune response can allow Mtb to persist, prolonged inflammation and loss of metabolic capacity can impair immune-cell function, promote tissue damage, and worsen disease. Consistent with these competing demands, pulmonary myeloid cells are highly heterogeneous during tuberculosis, and distinct macrophage populations can impose markedly different pressures on intracellular Mtb (Huang et al., 2018; Pisu et al., 2021). Thus, understanding tuberculosis requires defining not only which myeloid populations emerge during infection, but also the cellular programs that shape their functional states and how these programs change as infection progresses.

Mitochondria are increasingly recognized as central regulators of the macrophage response to Mtb (Patrick & Watson, 2021). In addition to their canonical role in ATP production through oxidative phosphorylation (OXPHOS), mitochondria shape innate immune responses by regulating cellular redox balance, generating mitochondrial reactive oxygen species (mtROS), releasing immunomodulatory mitochondrial molecules, and coordinating programmed cell death pathways (Bock & Tait, 2020; Chen et al., 2023). Several innate immune pathways that are activated during Mtb infection, including cGAS signaling (Watson et al., 2015), selective autophagy (Gutierrez et al., 2004; Watson et al., 2012), and apoptosis/necroptosis (Ding & Briken, 2026), are also triggered by mitochondrial damage and dysfunction, positioning mitochondria as a key node in the macrophage response to Mtb. Accordingly, Mtb has evolved ways to perturb host metabolism and mitochondrial integrity (Chen et al., 2006; Cumming et al., 2018; Lee et al., 2019; Pagan et al., 2022) and disruptions in mitochondrial homeostasis, including mutations in mitochondria-associated genes such as *LRRK2*, *POLG*, *PARK2*, *MFN2*, and *OPA1*, can significantly alter macrophage inflammatory and antimicrobial responses to mycobacterial infection (Mabry et al., 2025; Manzanillo et al., 2013; Silwal et al., 2021; Weindel et al., 2020; Weindel et al., 2022). Despite multiple lines of evidence pointing to mitochondria as critical nodes in controlling Mtb pathogenesis, we currently lack a framework to predict how a certain type of mitochondrial stress or mutation will impact Mtb infection outcomes.

Mitochondrial bioenergetics may represent an important component of this relationship. Macrophage activation is accompanied by extensive remodeling of glycolysis, the tricarboxylic acid cycle (TCA), and mitochondrial respiration (Kelly & O’Neill, 2015; Mehrotra et al., 2014). Unlike the simple Warburg shift from oxidative phosphorylation to glycolysis observed in macrophages stimulated with an innate agonist like LPS, Mtb infection elicits dynamic and context dependence shifts in macrophage bioenergetics (Cumming et al., 2018; Gideon et al., 2022; Howard & Khader, 2020). How these shifts play out over the course of a prolonged Mtb infection in the complex milieu of the lung and how these changes affect sustained antimycobacterial immunity *in vivo* remain poorly understood

Here, we define the temporal remodeling of mitochondrial respiratory programs across the pulmonary myeloid compartment during Mtb infection. By integrating single-cell transcriptional profiling with experimental disruption of mitochondrial electron transport in cultured macrophages, we describe shifts in OXPHOS transcriptional programming that are largely shared across diverse myeloid populations and associated with coordinated changes in metabolic and antimycobacterial pathways. We report that prolonged Mtb infection of macrophages is sufficient to induce similar reprogramming *ex vivo* and experimental impairment of Complex I recapitulates select transcriptional and functional features of the ETC^low^ state defined *in vivo*. Because similar transcriptional remodeling of OXPHOS accompanies Mtb seroconversion in tuberculosis household contacts, we conclude that mitochondrial respiratory remodeling is a shared feature of the myeloid response to Mtb in mouse models and humans. Collectively, these findings establish mitochondrial respiratory remodeling as a hallmark of the host response to Mtb and suggest that variation in mitochondrial energetic state may be a major determinant of tuberculosis disease outcomes.

## RESULTS

### Mtb disease progression dynamically remodels the immune cell landscape in the mouse lung

To define the cellular landscape of the immune response during *Mycobacterium tuberculosis* (Mtb) infection, we infected wild-type C57BL/6 mice using our low dose aerosol infection and performed single-cell RNA sequencing (scRNA-seq) on CD45+ cells (20,000 cells for uninfected and 10,000 cells for infected mice) enriched from the lungs of uninfected mice (n= 3, pooled) and mice at days 21 (n= 3, pooled) and 77 (n= 6, pooled) (**Fig. 1A**). Following Harmony integration, uniform manifold approximation and projection (UMAP) dimensional reduction, and unsupervised clustering, we identified 31 transcriptionally distinct populations representing major immune and stromal compartments of the lung. Using canonical marker genes, we identified T cells, NK cells, ILCs, B cells, monocytes, macrophages, dendritic cells, neutrophils, basophils, epithelial cells, endothelial cells, and fibroblasts (**Figs. S1A-D**). Consistent with activation of a robust immune response, we measure substantial changes in CD45+ cellular composition in Mtb infected vs. uninfected lungs. Notably, Mtb infection triggered an expansion of Cluster 1 (T cells 2), Cluster 17 (Macrophage 1) and Cluster 18 (Macrophage 2), and a depletion of Cluster 0 (T cells 1) and Cluster 19 (alveolar macrophages) (**Fig. S1C**). We also detected dynamic remodeling of the pulmonary immune landscape as Mtb disease progressed, with lymphocyte populations expanding at day 77 (most dramatically T cell 2, Tregs, and plasma cells) and alveolar macrophages and stromal cell populations proportionally reduced at day 77 (**Fig. S1D-E**). Similar shifts in lung cellular composition have been observed in murine and nonhuman primate TB, including increased representation of activated T cells and relative contraction of alveolar macrophages during disease progression (Duffy et al., 2026; Esaulova et al., 2021). These findings establish a comprehensive cellular atlas of the pulmonary immune response during Mtb infection and provide a framework for examining cell-type specific transcriptional changes as Mtb disease progresses in the mouse model.

**Figure 1.**
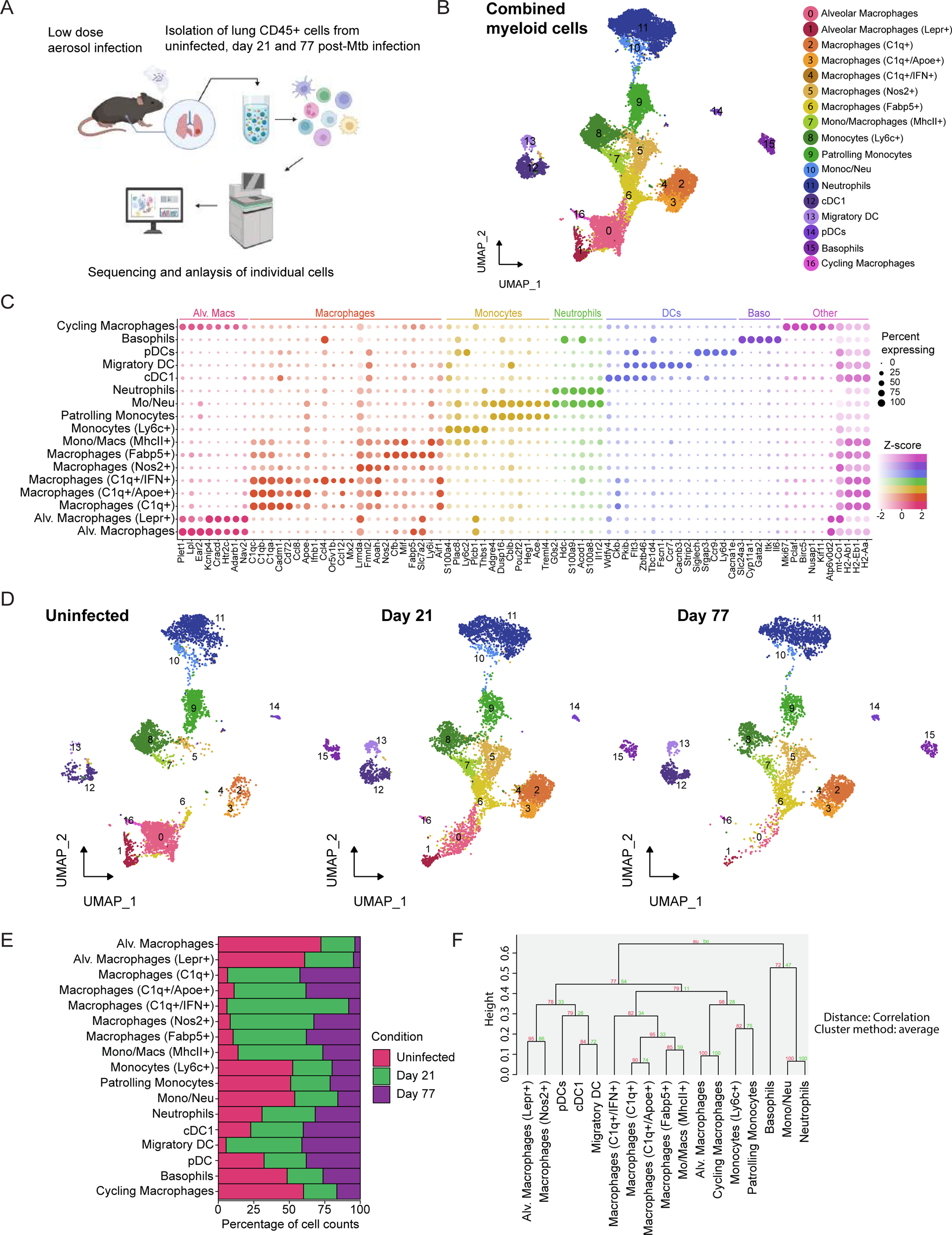
(A) Schematic representing the workflow used to generate single cell RNA sequencing data from CD45+ enriched lung cells for uninfected, 21 day post infection or 77 days post infection (B) UMAP visualization of aggregate data representing the myeloid immune landscape of Mtb infected mouse lung samples (uninfected, day 21 and day 77). (C) Dot plot showing representative marker genes for each annotated myeloid cell population. Dot size indicates percentage of cells expressing each gene and color intensity represents scaled gene expression (Z-score). Colors denote broad myeloid categories. (D) UMAP visualization of myeloid immune landscape of Mtb infected mouse lung samples of split by sample (uninfected, day 21 and day 77). (E) Bar plot showing relative abundance of cells across experimental conditions. Bars show percentage of cells contributed by each sample within each cell population. (F) Dendrogram showing the hierarchical clustering of myeloid populations based on the average expression of the 750 most variable genes using Pearsons correlation distance and average linking. Branch values indicate bootstrap support from 10,000 resamplings.

Since myeloid cells represent major cellular targets of Mtb infection and remain incompletely characterized at single-cell resolution during chronic disease, we next isolated and reanalyzed the myeloid compartment. Sub-clustering of myeloid cells identified 17 transcriptionally distinct populations consisting of alveolar macrophages, monocytes, dendritic cells, neutrophils, basophils, and multiple macrophage subsets (**Fig. 1B**). These included two alveolar macrophage populations, Ly6c+ inflammatory monocytes, patrolling monocytes, neutrophils, cDC1 cells, migratory dendritic cells, plasmacytoid dendritic cells, and several transcriptionally distinct macrophage populations defined by unique gene expression profiles. Cluster identities were assigned based on canonical marker gene expression (**Fig. 1C**): alveolar macrophage populations were characterized by expression of genes including *Plet1*, *Lpl*, and *Ear2*, whereas *C1q*-associated macrophage populations expressed high levels of *C1qa*, *C1qb*, and *C1qc*. Additional macrophage subsets were distinguished by expression of *Apoe, Ifnb1, Nos2*, *Fabp5*, or MHCII associated genes including *H2-Ab1*, *H2-Aa*, and *H2-Eb1*. Monocyte populations expressed *Ly6c2*, *Plac8*, and *Ace*, while neutrophils were defined by expression of *S100a8*, *S100a9*, *Acod1*, and *Il1r2*. Dendritic cell populations expressed canonical markers (e.g. *Flt3, Zbtb46, Fscn1, Ccr7, Siglech*, and *Ccr9*), supporting the identification of both conventional and plasmacytoid dendritic cell subsets. We identified two clusters that co-expressed canonical markers of macrophages or of neutrophils and monocytes labeled mono/mac or mo/neu respectively. These populations could represent transitional states, doublets or mixed lineage transcriptional profiles.

Projection of cells by infection status (uninfected, day 21, and day 77 post-infection) demonstrated substantial restructuring of the myeloid compartment following Mtb infection (**Fig. 1D**). While alveolar macrophages represented a dominant population in uninfected lungs, infected lungs contained an expanded diversity of macrophages and monocyte states. Several macrophage populations, including C1q+/Apoe+, C1q+/IFN+, Nos2+, Fabp5+, and MHCII+ monocyte/macrophage populations, were increasingly represented following infection and remained detectable throughout chronic disease. This expansion of heterogeneous infection-associated macrophage state is consistent with prior single cell studies demonstrating recruitment and diversification of monocyte derived and interstitial macrophage populations following pulmonary Mtb infection (Pisu et al., 2021; Zheng et al., 2025). In contrast, other populations (neutrophils and pDCs) exhibited comparatively stable representation across each timepoint. These findings indicate that Mtb infection is associated with the emergence and maintenance of diverse macrophage transcriptional states. Analysis of cluster frequencies further highlighted dynamic shifts in myeloid cell composition over the course of infection (**Fig. 1E**). Different macrophage, monocyte, dendritic cell, and granulocyte states dominated early and late infection, reflecting widespread diversification across myeloid lineages rather than alterations in a single cell type. To examine relationships among identified populations, we performed hierarchical clustering based on transcriptional similarity (**Fig. 1F**). This analysis grouped related macrophage populations while maintaining separation from monocyte, dendritic cell, neutrophil, and basophil populations, supporting the presence of multiple transcriptionally distinct myeloid states within the infected lung. Collectively, these findings reveal extensive heterogeneity within the pulmonary myeloid compartment and provide a comprehensive view of the transcriptional programs associated with myeloid responses during Mtb infection.

### Pulmonary myeloid cells undergo coordinated downregulation of OXPHOS and electron transport chain pathways during Mtb disease progression

Having established the cellular heterogeneity of the pulmonary myeloid compartment, we next sought to identify dominant transcriptional features shared across myeloid populations during disease progression. To accomplish this, we analyzed all myeloid cells as a single compartment and performed differential expression and gene set enrichment analyses (GSEA). Comparing day 21 vs. uninfected conditions, we observed a dramatic upregulation of pathways related to immune activation, interferon signaling, interferon response, antigen presentation/T cell activation, and complement (**Fig. S2A**). Similar pathway enrichment was seen in day 77 vs. uninfected conditions, with additional enrichment for genes linked to pathways related to lipid oxidation and fatty acid metabolism (downregulated) as well as B cell activation (**Fig. S2B**).

We next wanted to glean insight into how the myeloid compartment is transcriptionally remodeled as Mtb progresses from day 21 to day 77. Curiously, instead of revealing differences in cytokine signaling or antimicrobial defenses, comparison of transcriptional profiles between ay 21 and day 77 myeloid cells revealed widespread suppression of pathways associated with mitochondrial respiration and energy metabolism (**Fig. 2A**). Among the most significantly downregulated pathways were oxidative phosphorylation (OXPHOS), electron transport chain activity, aerobic respiration, ATP biosynthetic processes, and mitochondrial gene expression. Consistent with these findings, gene sets associated with oxidative phosphorylation from Gene Ontology Biological Process (GO:BP) and genes encoding oxidative phosphorylation machinery defined by Reactome exhibited strong negative enrichment in day 77 myeloid cells relative to day 21 (**Figs. 2B-D**).

**Figure 2.**
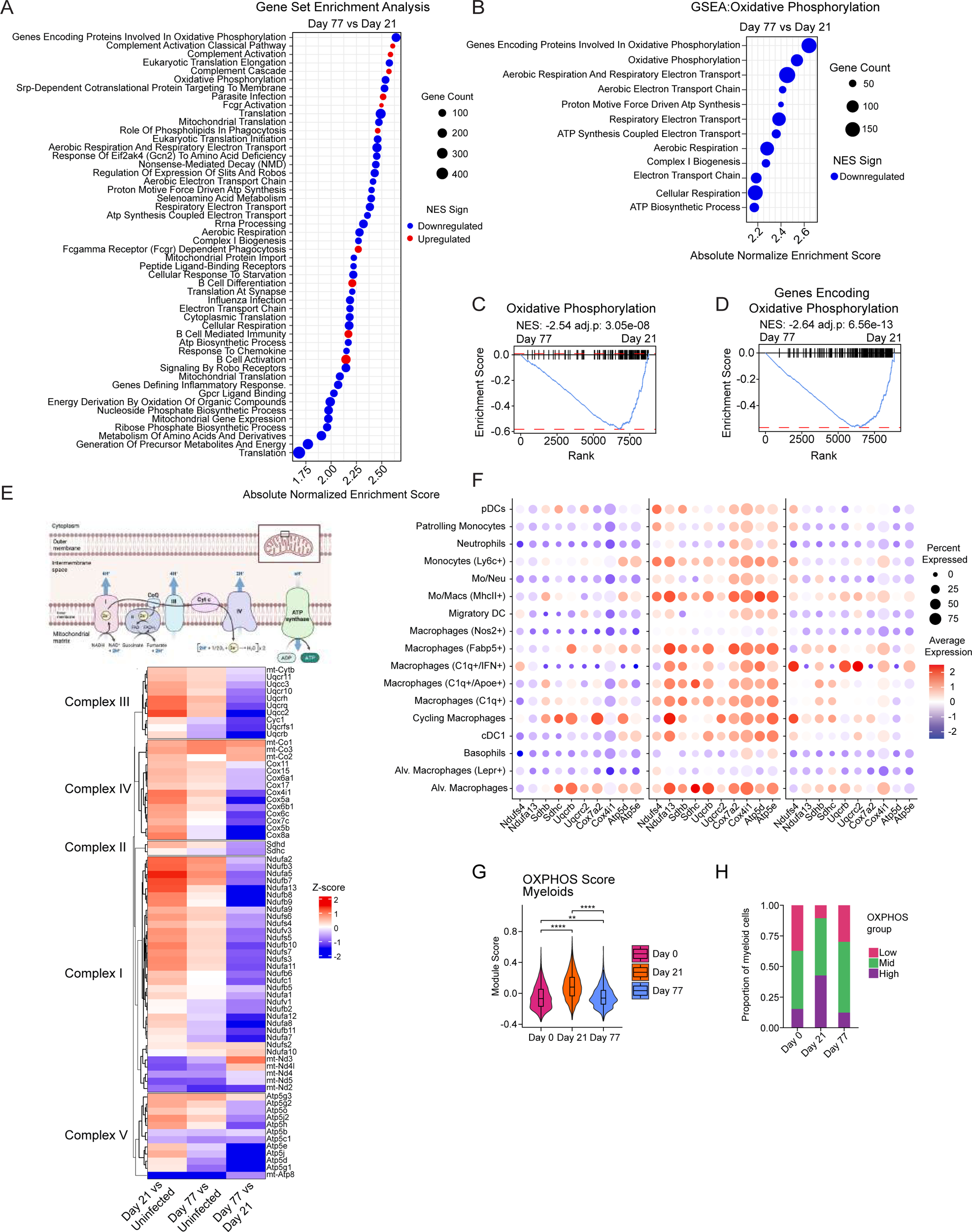
(A) Gene set enrichment analysis (GSEA) of aggregate myeloid transcriptional changes between day 77 and day 21, ranked by log2 fold change. Shown are pathways ordered by absolute normalized enrichment score NES, point size indicating gene set size, and color indicating direction; red is increasing enrichment and blue is decreasing enrichment at day77. (B) Selected OXPHOS and mitochondrial respiration pathways from the GSEA (C-D) GSEA enrichment plots the top OXPHOS pathway hits between day 77 and day 21 aggregate myeloid cells. NES and Benjamini-Hochberg adjust p values are indicated. (E) Heatmap of genes encoding OXPHOS complexes I-V, grouped by respiratory complex. Values represent z-scored log2 fold changes for the indicated comparisons. Associated graphics depict the OXPHOS complexes downregulated. (F) Dot plot of representative OXPHOS genes for each complex across the individual myeloid populations and infection time points. Dot size indicates percent expressed and color indicates average expression. (G) Violin plots of single cell OXPHOS modules scores across infection time points. Boxplots indicate median and interquartile range. Pairwise Wilcoxon rank-sum tests with Benjamini-Hochberg correction. (H) Bar plots showing the distribution of cells across low, intermediate and high OXPHOS states defined by module score quantiles.

To further examine the mitochondrial pathways underlying this enrichment pattern, we evaluated expression of genes encoding electron transport chain (ETC) Complexes I–V. Complexes I and II transfer electrons derived from NADH and succinate, respectively, to the ubiquinone, which feeds electrons to complex III, to cytochome C and complex IV. Electrons transfer through complexes I, III and IV is coupled with proton translocation across the inner mitochondrial membrane that generate the proton gradient that drives ATP synthesis by complex V (Vercellino & Sazanov, 2022). Consistent with our GSEA, many ETC associated transcripts were significantly elevated at day 21 post-infection but dramatically reduced by day 77 (**Fig. 2E, S2C**). This pattern was observed across genes encoding components of all respiratory chain complexes, including NADH dehydrogenase (Complex I), succinate dehydrogenase (Complex II), cytochrome bc_1_ complex (Complex III), cytochrome c oxidase (Complex IV), and ATP synthase (Complex V) subunits. Examination of representative ETC genes demonstrated a similar trend across multiple myeloid populations, indicating that loss of OXPHOS-associated transcriptional programs was not restricted to a single cell type (**Fig. 2F**).

To determine whether these changes reflect broad alterations across different cell types in the myeloid compartment, we calculated oxidative phosphorylation module scores for individual populations. The overall myeloid compartment exhibited a transient increase in OXPHOS score at Day 21 followed by a significant decline at Day 77 (**Fig. 2G-H**). Although basal OXPHOS scores differed among myeloid populations, with differentiated macrophage subsets including alveolar, Fabp5+, and C1q+ macrophages generally exhibiting higher OXPHOS scores than migratory monocytes, neutrophils, basophils, and pDCs, nearly all populations displayed the same temporal pattern of increased OXPHOS at Day 21 followed by decline at Day 77 (**Fig. S2D**). Together, these findings demonstrate that OXPHOS undergoes coordinated temporal remodeling across the pulmonary myeloid compartment during Mtb infection. Importantly, reduced expression of OXPHOS genes did not occur in the context of global transcriptional suppression, as numerous pathways demonstrated dramatic upregulation in the same cell populations (**Fig. S2A-B**). These observations suggest that chronic infection is characterized not by a loss of overall transcriptional activity in myeloid cells, but rather by selective remodeling of metabolic programs, including progressive suppression of genes associated with mitochondrial respiration and oxidative phosphorylation.

### Declining OXPHOS is associated with broad remodeling of metabolic and immune programs in pulmonary myeloid cells

Having established that OXPHOS gene expression declines as Mtb infection progresses, we next asked whether differences in OXPHOS were linked to broader changes in myeloid cell gene expression that could hint at altered function. Within each myeloid population, we correlated the expression of every gene with OXPHOS module scores (**Fig. 2G**) and ranked them accordingly. We then performed Gene Set Enrichment Analysis (GSEA) independently for each population. Pathway Normalized enrichment scores were averaged across the associated myeloid populations to identify programs that are consistently associated with OXPHOS with dot size representing the number of myeloid populations associated with the pathway (full list **Fig. S3A**, curated list **Fig. 3A**). Pathways related to mitochondrial respiration, protein synthesis, RNA surveillance, antigen processing, interferon responses, reactive oxygen species, and glycolysis were significantly enriched for genes that positively correlate with OXPHOS score (**Fig. 3A and S3A, red**). In contrast, inflammatory response, in particular TGF-β associated genes, were enriched among genes negatively correlated with OXPHOS (**Fig. 3A and S3A-B, blue**).

**Figure 3.**
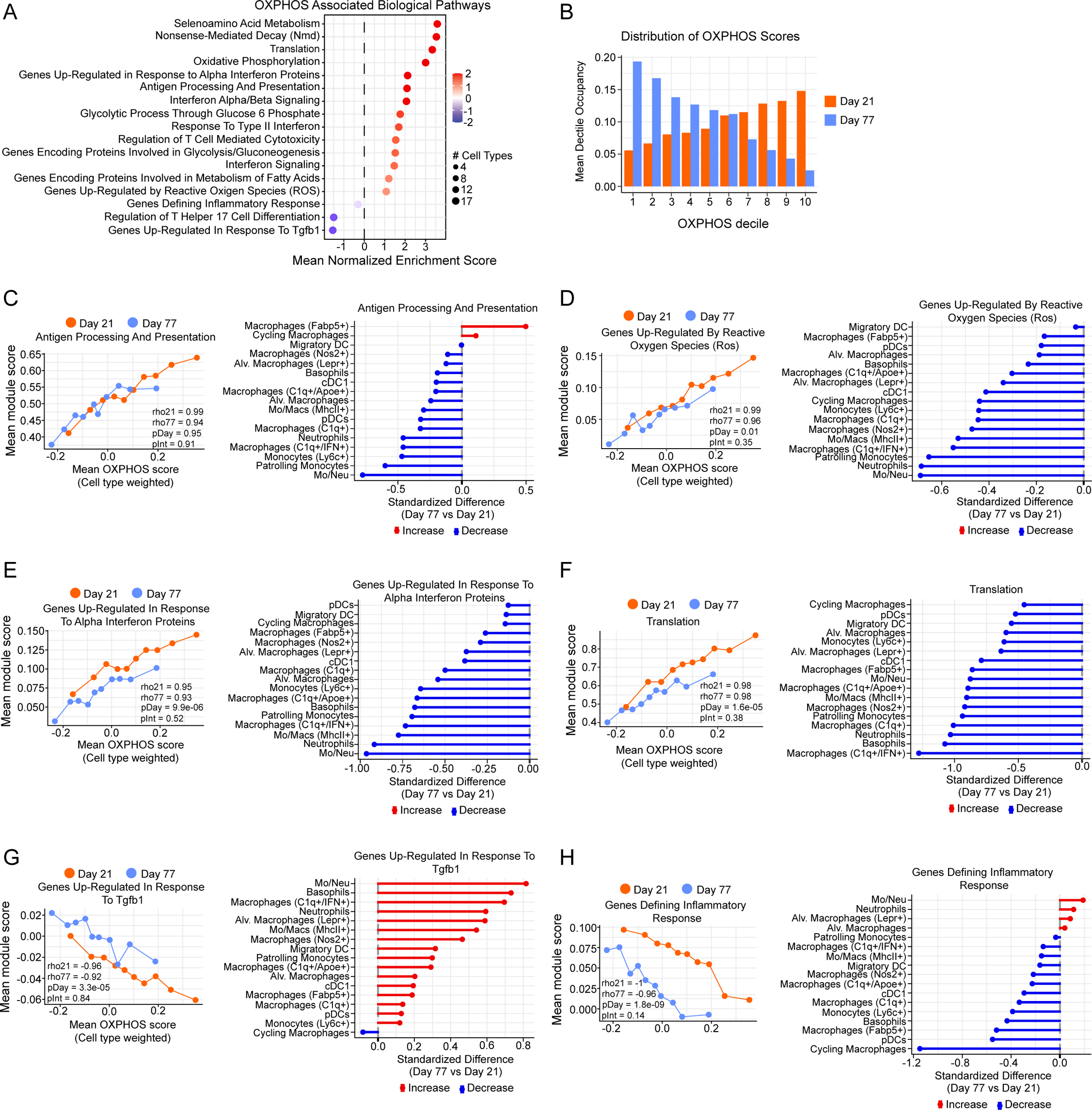
(A) GSEA of selected pathways. Genes ranked by their Spearman correlation with OXPHOS module score within individual myeloid populations. Selected pathways are significantly enriched in at least two myeloid populations are summarized by mean normalized enrichment score (NES) and dot size represents the number of cell types the pathway was significant. (B) Distribution of Day 21 and Day 77 cells across OXPHOS score dectiles. Mean dectile occupancy represents the average proportion of cells within each dectile across individual myeloid populations. (C-H) Line plots show the relationship between OXPHOS state and module scores for indicated pathways at day 21 and 77. Values represent cell type weighted summaries. Spearman correlations were calculated separately for each day, and linear models test day and day by OXPHOS interactions. Accompanying the lines plots are lollipop plots show the standardized differences in pathway module scores between day 77 and day 21 across individual populations. Cohen’s d was calculated as day 77 relative to day 21; positive values (red) indicate higher module scores at day 77 while negative values (blue) indicate lower module scores.

We next asked if the progressive loss of OXPHOS in the myeloid compartment was accompanied by a redistribution of different cell subtypes towards lower OXPHOS states. By ranking Day 21 and Day 77 cells together and dividing them into OXPHOS deciles (accounting for differences in baseline OXPHOS between myeloid populations), we observed that Day 21 cells were more frequently distributed among higher OXPHOS deciles, whereas Day 77 cells shifted toward lower OXPHOS deciles (**Fig. 3B**). Thus, chronic infection not only lowered the average OXPHOS score but also shifted the distribution of cells toward lower OXPHOS deciles, reflecting a population wide redistribution along the OXPHOS continuum.

Next, we asked how the OXPHOS-associated pathways identified by GSEA (**Fig. 3A**) varied between cells with relatively high and low OXPHOS states and whether these relationships were maintained between Day 21 and Day 77. Unlike the redistribution analysis (**Fig. 3B**), OXPHOS deciles were defined separately within each cell type and time point to allow pathway activity and OXPHOS state to be evaluated independently. Consensus leading-edge genes across the myeloid compartment from the GSEA were used to calculate pathway module scores. OXPHOS and pathway module score medians were calculated within each cell type and decile then averaged across cell types, giving each population equal contribution to the resulting trajectory analysis. As expected, OXPHOS module score increased across the OXPHOS deciles at both time points (**Fig. S3B**).

Notably, we observed that antigen presentation (**Fig. 3C**), reactive oxygen species (**Fig. 3D**), interferon response (**Fig. 3E**), and translation (**Fig. 3F**) were positively correlated with OXPHOS at both Day 21 and Day 77. Although pathway activity varied based on OXPHOS state, no significant day effects or day-OXPHOS interactions were apparent. We also quantified the extent to which these pathways were remodeled during Mtb infection (magnitude and direction) by measuring effect size between day 77 vs day 21. Despite a role for dendritic cells, alveolar macrophages and neutrophils in antigen presentation, lipid handling, and glycolytic metabolism (Del Prete et al., 2023; Grudzinska et al., 2023; Wculek et al., 2023), these pathways were down in these populations (**Fig. 3C-E**, lollipop graphs). In contrast, Fabp5+ macrophages (a population of lipid-rich macrophages) and cycling (proliferating) macrophages displayed increased antigen processing and presentation, suggesting some plasticity in how certain subtypes of myeloid cells respond to low OXPHOS (**Fig. 3C**). TGFB associated genes displayed an inverse relationship with OXPHOS, (low expression when OXPHOS is high, high with OXPHOS is low) (**Fig. 3G**) but showed Day effects as well (i.e. their overall activity differed between Day 21 and Day 77). Interestingly, although genes involved in the inflammatory response displayed an inverse relation to OXPHOS (i.e. inflammatory gene expression is higher in cells with low OXPHOS gene expression), this effect was largely eclipsed by a significant Day effect, whereby inflammation was overall much higher at Day 21 compared to Day 77 (**Fig. 3H**). These results suggest that although there may be a relationship between OXPHOS decline and inflammation, additional infection stage-dependent inputs also influence inflammatory gene expression over the course of Mtb infection *in vivo*. Collectively, these analyses demonstrate that low OXPHOS myeloid cells experience remodeling of other immune-relevant pathways in the lungs of Mtb-infected mice and begin to suggest that OXPHOS decline could be driving these changes.

### Progressive Electron Transport Chain Dysfunction Underlies Metabolic Suppression During Mtb Infection

ETC low myeloid cell populations isolated from the lungs of Mtb-infected mice are exposed not only to Mtb-derived PAMPs and DAMPs but also to adaptive immune cells, hypoxia, and circulating cytokines. To determine if Mtb infection is sufficient to induce OXPHOS transcriptional reprogramming in macrophages, we took a reductionist *ex vivo* approach. Briefly, we infected bone marrow derived macrophages (BMDMs) from wild-type C57BL6 mice with Mtb (Erdman) MOI = 1 and quantified differences in transcript abundance by 3’ end counting RNA-seq over an extended infection time course (day 1, 3, and 5 post-infection). Consistent with our scRNA-seq of myeloid cell populations, we observed a modest, but global, downregulation of genes encoding all five components of the electron transport chain in macrophages infected with Mtb for 3 and 5 days (relative to Day 1) (**Fig. 4A-B**). ETC components were also downregulated at the protein level by immunoblot, using antibodies directed against representative components of Complex I (NDUFS4) (**Fig. 4C**), Complex II (SDHB) (**Fig. S4A,B**), Complex IV (COXIV) (**Fig. 4B**) and Complex V (**Fig. S4A,D**). No change in the Complex III protein UQCRC2 was detectable (**Fig. S4A,C**). Further mirroring our scRNA-seq data, gene set enrichment analysis of Day 3 and 5 RNA measured enrichment for downregulated genes in pathways related to oxidative phosphorylation, glycolysis, interferon alpha response, and fatty acid metabolism, and increased expression of genes involved in TGF-β signaling (**Fig. 4D**). These data suggest that Mtb infection of macrophages is sufficient to trigger the transcriptional reprogramming we observed in the lungs of Mtb infected mice. Interestingly, chronic exposure of BMDMs to PAM3CSK4 (a TLR2 agonist) (**Fig. 4E-F**) or IL-1β (an inflammatory cytokine frequently hyperinduced by cells experiencing mitochondrial dysfunction (**Fig. 4G-H**) (Shimada et al., 2012; Weindel et al., 2022; Xian et al., 2022)) was sufficient to downregulate Complex I transcript abundance (*Ndufa1* and *Ndufc2*), suggesting that integration of multiple innate sensing events likely drives ETC downregulation in macrophages during Mtb infection.

**Figure 4.**
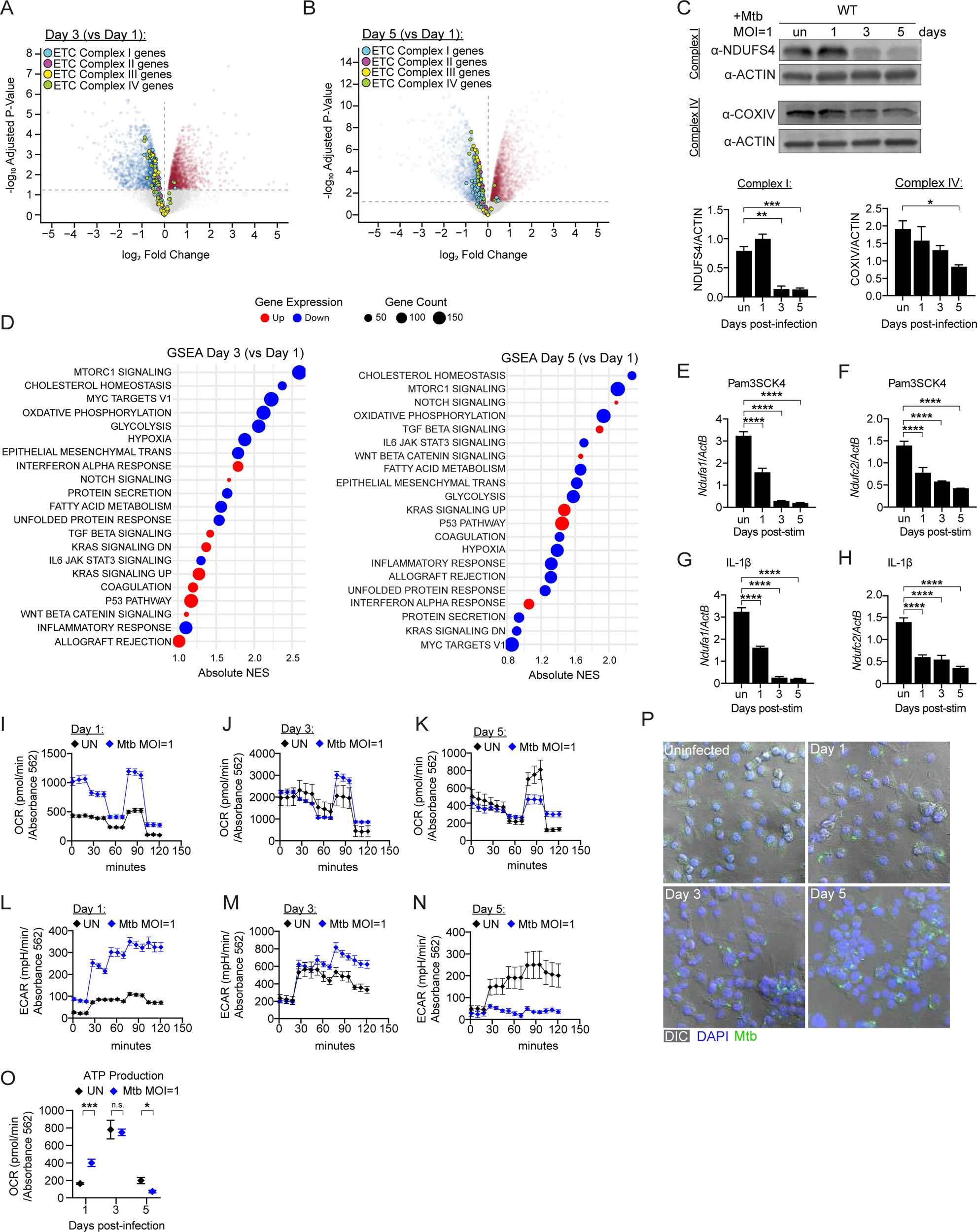
(A) Volcano plot of differentially expressed genes in Mtb-infected (MOI=1) BMDMs at Day 3 compared to Day 1. ETC complex I, II, III, IV genes were annotated based on the MitoCarta3.0. Upregulated genes in red. Downregulated genes in blue. (p≤0.05) (B) As in A, but in Mtb-infected (MOI=1) BMDMs at Day 5 compared to Day 1. Upregulated genes in red. Downregulated genes in blue. (p≤0.05) (C) Immunoblot analysis of Ndufs4 and CoxIV in uninfected and Mtb-infected (MOI=1) BMDMs at 1, 3, and 5 days. Actin was used as a loading control. Quantification of Ndufs4 and CoxIV protein levels normalized to Actin. n=3 (D) GSEA pathway dot plots showing the top upregulated and downregulated Hallmark pathways ranked in Mtb-infected BMDMs (MOI=1). Day 3 versus Day 1 is shown on the left, and day 5 vs day 1 is shown on the right. Genome-wide ranking vector ordered by log_2_Fold Change for each comparison. Pathways are ordered on the y-axis by Normalized Enrichment Score (NES). Dot size scales proportionally with the number of core enrichment genes contributing to the pathway. Color specifies upregulation (red) or downregulation (blue). Statistical filtering was controlled using Benjamini-Hochberg adjusted p-values. (E) qRT-PCR gene expression of *Ndufa1* in untreated and 20 ng/mL Pam3CSK4 treated BMDMs for 1, 3, and 5 days. Gene expression was normalized to *Actb*. (F) As in E, but measuring *Ndufc2*. (G) qRT-PCR gene expression of *Ndufa1* in untreated and 1 ng/mL IL-1β treated BMDMs for 1, 3, and 5 days. Gene expression was normalized to *Actb*. (H) As in G, but measuring *Ndufc2*. (I) Oxygen consumption rate (OCR) measured by Agilent Seahorse Metabolic Analyzer in uninfected and Mtb-infected (MOI=1) BMDMs at Day 1. (J) As in I, but OCR at Day 3 post-Mtb infection. (K) As in I, but OCR at Day 5 post-Mtb infection. (L) Extracellular acidification rate (ECAR) measured by Agilent Seahorse Metabolic Analyzer in uninfected and Mtb-infected (MOI=1) BMDMs at Day 1. (M) As in L, but OCR at Day 3 post-Mtb infection. (N) As in L, but OCR at Day 5 post-Mtb infection. (O) ATP production as measured by Agilent Mito Stress test in uninfected and Mtb-infected (MOI=1) BMDMs at 1, 3, 5 days post-Mtb infection. P) Representative DIC and IF images of uninfected and Mtb-infected (MOI=1) BMDMs at 1, 3, 5 days. IF microscopy Mtb (green, anti-Mtb) and nucleus (blue, DAPI). Statistical analysis: *p < 0.05, **p < 0.01, ***p < 0.001, ****p < 0.0001. Statistical differences were determined for (C, E-H) using one-way ANOVA with Tukey’s post-test and (O) using two-way ANOVA with Sidak’s post-test.

To begin to link altered ETC gene expression with OXPHOS disruption, we set out to measure mitochondrial output at Mtb-infection timepoints when ETC gene expression is depressed. We infected BMDMs with Mtb Erdman (MOI=1) and measured ECAR (glycolysis) and OCR (OXPHOS) at 1, 3, and 5 days post-infection using the Agilent Seahorse Metabolic Analyzer. Over the course of 5 days, Mtb-infected BMDMs displayed a biphasic metabolic trajectory, with increased OCR and ECAR at day 1 post-infection (**Fig. 4I, L**), followed by a decline in basal respiration, maximal respiration, spare respiratory capacity, and ATP production by day 5 post-infection (**Fig. 4J-K, S4E**). Somewhat unexpectedly, this decrease in OCR was not accompanied by a compensatory increase in glycolysis, as maximal glycolytic capacity and glycolytic reserve were both significantly decreased at Day 5 post-infection (**Fig. 4M-N, S4F)**. Consistent with decreased OXPHOS and glycolysis, an overall decrease in cellular ATP was measured at Day 5 post-Mtb (MOI=1) (**Fig. 4O**). To ensure that downregulation of ETC genes, loss of ATP, and altered metabolic output did not result from unhealthy/dying cells at late infection timepoints, we changed media daily and monitored monolayers and total protein collected for all our 5-day experiments. Monolayers remained healthy and intact over the course of infection (**Fig. 4P**) and protein levels were consistent between uninfected and Mtb-infected cells at each timepoint (**Fig. 4C, S4G**).

Because these results stand in contrast to the “bioenergetic senescence” phenotype previously ascribed to Mtb at 24h post-infection (Cumming et al., 2018), we also performed infections at MOI=5 and 10, to more accurately recapitulate earlier studies. We confirmed that at higher MOIs, this entry into “bioenergetic paralysis” is hastened, with dampened maximal respiration and spare respiratory capacity measured at 24h post-Mtb at MOI=5, and further exacerbation seen at MOI=10 (**Fig. S4H**). Total protein levels were again comparable at each MOI (**Fig. SI**). Based on these observations, we predict that a certain threshold of stress needs to be met to trigger bioenergetic senescence in macrophages and that this happens faster at higher MOIs. Finally, to examine how limited metabolic output might influence other aspects of the macrophage response to Mtb, we leveraged our 3’ end counting RNA-seq (**Fig. S4J**). Using Reactome GSEA, we identified MHC Class II Antigen Presentation (*Cd74*, *H2-Eb1*, *H2-DMb1*, *Ctsh*, and *Ctsc*) (**Fig. S4K**), Interferon signaling (*Irf4*, *Irf8*, *Socs3*, *Mavs*, and *H2-Eb1*) (**Fig. S4L**), and phagocytosis (*Fcgr1*, *Fcgr2b*, *Fcgr4*, *Syk*, *Prkcd*, and *Prkce*) (**Fig. S4M**) among the most downregulated pathways in Mtb-infected macrophages at Day 5 post-infection. Vesicle-mediated transport and membrane trafficking genes were upregulated (*Lrp1*, *Cd36*, *App*, *Snap23*, *Stxbp3*, *Tubb4b*, and *Kif5b*) (**Fig. S4N**). These data start to suggest that functional differences in innate immune pathways accompany changes to ETC gene expression and depressed mitochondrial respiration in Mtb-infected macrophages.

### Loss of Complex I function in macrophages creates a permissive niche for Mtb

Having observed that (1) prolonged infection of macrophages with Mtb downregulates macrophage OXPHOS and (2) transcriptional reprogramming of antimycobacterial immune pathways correlates with ETC downregulation, we next wanted to more directly link ETC dysfunction to the macrophage response to Mtb. To this end, we generated immortalized BMDMs (iBMDMs) stably expressing an shRNA targeting NADH:ubiquinone oxidoreductase subunit S4 (*Ndufs4*), a well-characterized component of mitochondrial Complex I (**Fig. 5A**). As expected, *Ndufs4*-deficient macrophages exhibited impaired mitochondrial respiration, including reductions in basal respiration, maximal respiration, spare respiratory capacity, and ATP linked respiration (**Fig. 5B** and **S5A**). Alterations in glycolytic capacity were also observed in *Ndufs4* KDs (**Fig. 5C** and **Fig. S5B**), supporting our use of these cells as representative of the ETC^low^/glycolysis^low^ myeloid populations we identified *in vivo* (**Fig. 3**).

**Figure 5.**
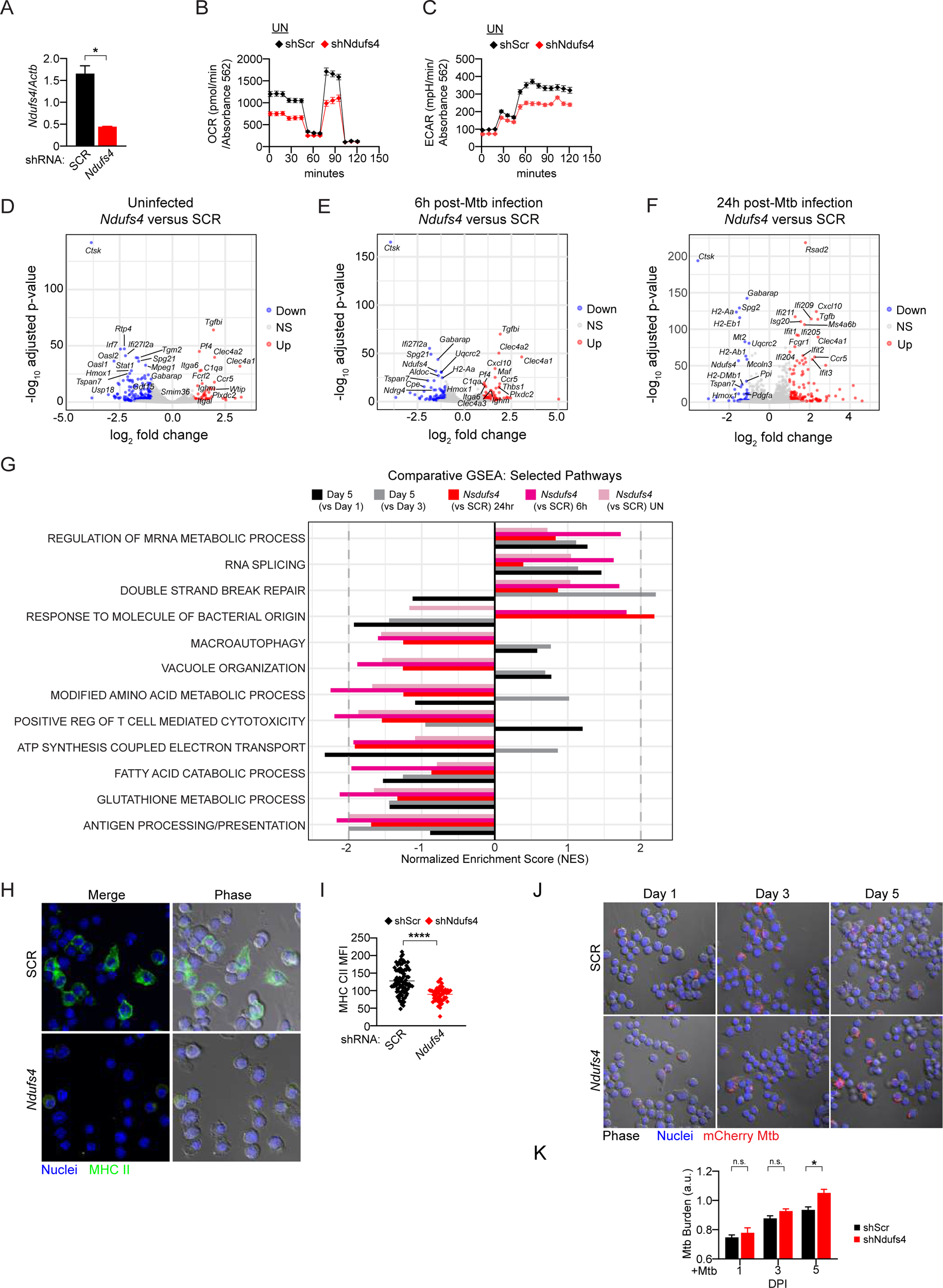
(A) qRT-PCR gene expression of *Ndufs4* in untreated SCR KD and *Ndufs4* KD iBMDMs. Gene expression was normalized to *Actin*. (B) Oxygen consumption rate (OCR) measured by Agilent Seahorse Metabolic Analyzer in untreated SCR KD and *Ndufs4* KD iBMDMs. (C) Extracellular acidification rate (ECAR) measured by Agilent Seahorse Metabolic Analyzer in untreated SCR KD and *Ndufs4* KD iBMDMs. (D) Volcano plots differentially expressed genes in uninfected SCR KD and *Ndufs4* KD iBMDMs. The x-axis plots log_2_Fold Change values against -log_10_-transformed adjusted p-values on the y-axis. Statistical significance for genes was defined by an adjusted p-value < 0.05 combined with absolute log_2_Fold change > 1. Red indicates upregulation, blue indicates downregulation, and grey indicates non-significance. The top differentially expressed genes are annotated. (E) As in D, but at 6h post-Mtb infection (MOI=5) SCR KD and *Ndufs4* KD iBMDMs. (F) As in D, but at 24h post-Mtb infection (MOI=5) SCR KD and *Ndufs4* KD iBMDMs. (G) Grouped bar graph displaying NES for select GO:BP pathways across multiple experimental comparisons. Transcripts ranked continuously by log_2_Fold Change from each corresponding DESeq2 contrast. Overlapping cohorts are color-coded by experimental group: Day 5 vs Day 1 (Mtb-infected MOI=1) (black), Day 5 vs Day 3 (Mtb-infected MOI=1) (grey), *Ndufs4* vs SCR uninfected control (light pink), *Ndufs4* vs SCR 6h post-Mtb infection (hot pink) and *Ndufs4* vs SCR 24h post-Mtb infection (red). Pathways are ordered on the y-axis by Normalized Enrichment Score (NES). (H) Immunofluorescence microscopy of untreated SCR KD and *Ndufs4* KD iBMDMs. Nucleus (DAPI, blue) and MHC CII (anti-MHC CII, green). (Scale bar) (I) Quantification of IF in H. Mean fluorescence intensity (MFI) defined by DIC/BF-based cell mask and calculated within a specified region of interest (a.u.). (J) Immunofluorescence microscopy of uninfected and mCherry Mtb-infected (MOI=1) SCR KD and *Ndufs4* KD iBMDMs at 1, 3, and 5 days. Nucleus (DAPI, blue) and Mtb (mCherry, red). (Scale bar) K) Quantification of IF in J. Mtb bacilli burden was quantified via bacilli segmentation and mask construction to measure subsequent bacilli area on a per cell basis. Bacilli area (a.u.) normalized to t=0h mCherry Mtb-infected SCR KD and *Ndufs4* KD iBMDMs. Statistical analysis: *p < 0.05, **p < 0.01, ***p < 0.001, ****p < 0.0001. Statistical differences were determined for (A) using a two-tailed Student’s unpaired t test, (I) using an unpaired Welch’s t test, and (K) using multiple two-tailed Student’s unpaired t tests.

3’end counting RNA-seq of *Ndufs4* KDs vs. SCR controls revealed substantial transcriptional remodeling in both uninfected and Mtb-infected cells (**Fig. 5D-F**). In uninfected *Ndufs4* KDs macrophages, we noted dampened expression of type I IFN genes (*Irf7*, *Oasl1*, *Oasl2*, *Stat1*, *Ifi27l2a*) relative to SCR (**Fig. 5D, S5C**). This phenotype was less pronounced at 6h post-Mtb infection, likely representing a transitional state as previously downregulated inflammatory and type I IFN genes begin to be induced (**Fig. 5E, S5D**). At 24h post-infection, *Ndufs4* KDs exhibited a dramatic failure to induce genes related to MHC Class II presentation, with *H2-Aa*, *H2-Ab1*, *H2-Eb1*, and *H2-DMb1* all downregulated compared to SCR (**Fig. 5F, S5D-E**). This finding nicely aligns with previous work, demonstrating that Complex I is required for IFN-γ-induced expression of MHCII genes and downstream T cell activation (Kiritsy et al., 2021). At 24h post-Mtb infection, we observed a failure to induce stress responsive genes like *Hmox1* in *Ndufs4* KD cells, alongside hyperinduction of type I IFN genes, hinting at altered NRF2-function in Complex I-deficient cells (**Fig. 5F**). Interestingly, significant downregulation of cathepsin K (*Ctsk*), a lysosomal cysteine protease, was seen in *Ndufs4* KDs in all conditions (**Fig. 5D-F**). Collectively, these in vitro analyses support our claim that loss of mitochondrial respiratory capacity induces broad transcriptome changes in macrophages.

Finally, to gain insight into how cells genetically engineered to downregulated Complex I compare to wild-type cells undergoing this transition in response to Mtb infection, we performed pathway enrichment analysis comparing wild-type macrophages infected with Mtb at 3- and 5-day post-infection with *Ndufs4* KD cells at 0, 6, and 24h post-Mtb (**Fig. 5G**). Pathway enrichment was largely shared across all conditions, supporting the idea that Day 5 post-Mtb macrophages phenocopy *Ndufs4* KD macrophages. Some notable differences were seen in macroautophagy, vacuole organization, and amino acid metabolism, which were upregulated in wild-type macrophages infected with Mtb (grey/black) but downregulated in *Ndufs4* KDs (pink/red), hinting at a role for Complex I in activation of these pathways (**Fig. 5G**). Interestingly, pathways related to antigen presentation, ATP synthesis coupled electron transport, glutathione metabolism, and fatty acid catabolism were similarly downregulated in wild-type and Ndufs4 KD macrophages, but with vastly different kinetics (**Fig. 5G**), suggesting that loss of Complex I may hasten the transcriptional reprogramming that we see at day 3 and 5 post-Mtb infection of wild-type macrophages.

We then set out to explore whether transcriptional changes in *Ndufs4* KD macrophages functionally impacted innate immune responses. Having observed downregulation of macroautophagy and vacuolar organization genes in *Ndufs4* KD macrophages infected with Mtb (**Fig. 5G**), we asked whether these cells could effectively restrict Mtb replication, using mCherry signal as a proxy for bacterial burden. At day 5 post-infection, mCherry MFI was significantly higher in *Ndfus4* KD iBMDMs compared to SCR controls (**Fig. 5H-I**), implicating Complex I function in cell-intrinsic control of Mtb. Likewise, dampened expression of genes involved in antigen presentation emerged as a recurring feature of both ETC^low^ myeloid cell and *Ndufs4* KD datasets (**Fig. 5G**). To ask whether these transcriptional changes affect MHC Class II at a functional level, we performed immunofluorescence microscopy using antibodies designed to detect extracellular MHCII in *Ndufs4* and SCR iBMDMs and quantified MFI per cell. In resting cells, we measured significantly less MHCII on *Ndufs4* KD cells compared to controls (**Fig. 5J-K**). This finding, together with our transcriptional analyses, support a role for Complex I in maintaining antigen presentation in macrophages and is consistent with a model in which loss of ETC function disrupts macrophage effector programs during chronic Mtb infection in mice.

### Downregulation of ETC genes correlates with Interferon-Gamma Release Assay (IGRA) seroconversion in humans

To determine whether the decline in OXPHOS associated programs observed in mouse myeloid cells was also detectable in humans exposed to Mtb, we analyzed scRNA-seq data generated from close contacts of culture-confirmed pulmonary TB cases from a RePORT-Brazil cohort. Peripheral blood mononuclear cells (PBMCs) collected from 45 Mtb exposed household contacts were stratified by interferon gamma release assay (IGRA) status, distinguishing individuals with detectable Mtb-specific T cells responses from those without an IGRA response (Pena Avila et al., 2025)(**Fig. S6A**). We first asked whether myeloid cells from IGRA-positive individuals exhibited similar transcriptional changes to those identified in pulmonary myeloid populations from chronically infected mice. Gene set enrichment analysis of differentially expressed genes identified broad suppression of pathways associated with mitochondrial respiration and protein biosynthesis in IGRA-positive myeloid cells. Oxidative phosphorylation, electron transport chain, ATP synthesis-coupled electron transport, proton motive force-driven ATP synthesis, ATP biosynthetic processes, cellular respiration, translation, nonsense-mediated decay, and rRNA processing were de-enriched, whereas pathways involved in vesicle organization and endomembrane system organization were positively enriched (**Fig. 6A**). We next asked whether these metabolic programs were conserved across individual myeloid populations. Although the magnitude of pathway enrichment differed among subsets, reduced oxidative phosphorylation, electron transport chain activity, ATP biosynthetic processes, translation, and nonsense-mediated decay were detected across multiple circulating myeloid populations, while vesicle and endomembrane organization exhibited the opposite trend (**Fig. 6B**). Classical and activated monocytes showed the strongest reduction for OXPHOS associated gene sets. cDCs were less affected, with significant reductions limited to the Respiratory Electron Transport gene set. Intermediate monocytes did not show significant reductions in OXPHOS associated pathways, but retained a decrease in translation and RNA processing, whereas pDCs showed little to no enrichment (**Fig. 6B**). These observations parallel our mouse scRNA seq data in which we see loss of OXPHOS programs broadly across the myeloid compartment with monocyte and macrophage populations showing the most pronounced reductions. Consistent with pathway enrichment, genes encoding multiple electron transport chain complexes were broadly reduced in classical monocytes from IGRA+ individuals (i.e. *NDUF* genes encoding Complex I subunits, *SDH* genes of Complex II, *COX* genes of Complex IV, and *ATP5* genes of Complex V/ATP synthase) (**Fig. 6C**). Expression of Complex III genes were not significantly changed. These findings indicate that suppression of OXPHOS associated transcriptional programs is associated with IGRA+ seroconversion following Mtb exposure in humans and argue that remodeling mitochondrial respiratory programs represents a conserved feature of the myeloid response to Mtb across species.

**Figure 6.**
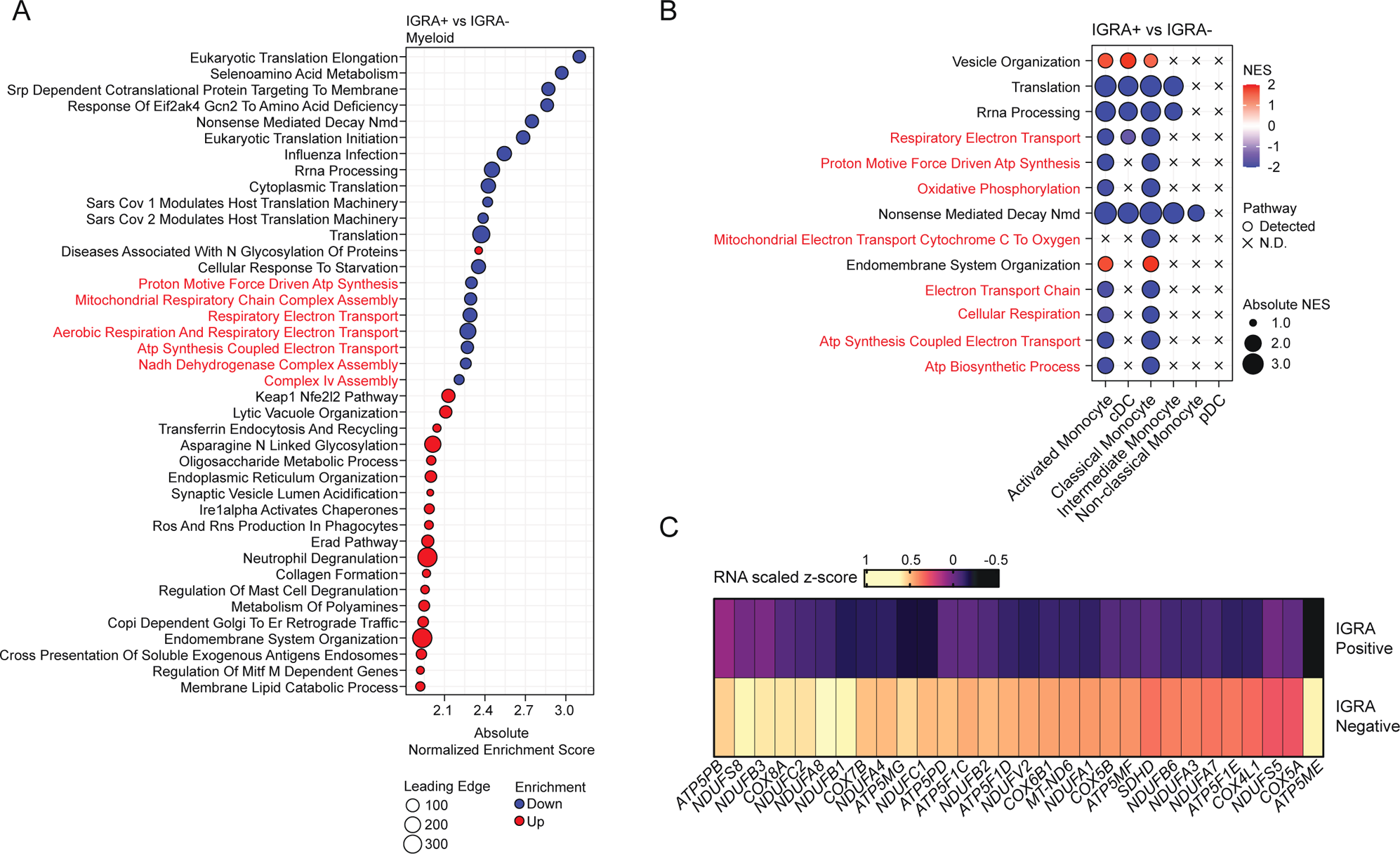
(A) GSEA of aggregate myeloid compartment of PBMCs comparing IGRA+ vs IGRA negative individuals. The 20 most positively and negatively enriched GO Biological Process and Reactome pathways are shown. Point positive indicates absolute normalized enrichment score (NES) color indicates increasing (red) or decreasing (blue) enrichments and point size indicates gene set size. (B) GSEA of selected mitochondrial respiratory and relate pathways across individual myeloid populations. Colors indicates NES and point size indicates absolute NES. N.D., not detected in the GSEA result. (C) Heatmap of genes encoding OXPHOS complexes I-V (complex III was not represented). Values are of gene scaled expression (z score).

## DISCUSSION

Little is known about how the immunometabolic landscape of the lung evolves over the course of tuberculosis infection. Here, using a mouse model of Mtb pathogenesis, we show that prolonged Mtb infection is associated with a broad metabolic transition across the pulmonary myeloid compartment, in which diverse myeloid populations converge toward a lower mitochondrial respiratory state. Our findings identify mitochondrial respiratory capacity as a shared feature of myeloid-cell state during tuberculosis and suggest that metabolic remodeling is dynamic rather than pre-determined by myeloid precursor/lineage.

A central finding of this study is the biphasic nature of metabolic transcriptional reprogramming. Following an initial increase in both OXPHOS and glycolysis at 24h, Mtb-infected macrophages *ex vivo* undergo an energetic collapse (Day 3 and 5), characterized by a decrease in OXPHOS and glycolysis, and loss of mitochondrial transcripts and protein (**Fig. 4**). This response is largely mirrored in pulmonary myeloid populations from Mtb-infected mice, whereby glycolysis and ETC-related genes are globally upregulated at day 21 post-infection and downregulated at day 77. In conventional low-dose aerosol C57BL/6 models, Mtb actively replicates during the first 2–3 weeks of infection before adaptive responses kick in and bacterial burdens stabilize (Kramnik & Beamer, 2016; Munoz-Elias et al., 2005). Thus, at day 21 post-infection, we would predict that new myeloid cells continue to infiltrate the lung, providing Mtb new replicative niches. As these macrophages encounter Mtb for the first time, they will need to upregulate multiple energy-generating pathways to mount inflammatory and antimicrobial responses (i.e. akin to macrophages *ex vivo* 24h post-infection). At later timepoints (day 77 *in vivo* or day 5 post-infection *in vitro*), pulmonary myeloid cells fail to maintain this high energy-generating state (**Fig. 2**). These data align with earlier studies of cultured human macrophages, wherein Mtb was shown to induce a state of decelerated bioenergetic metabolism characterized by decelerated flux through glycolysis and TCA and increased dependency on exogenous fatty acids (Cumming et al., 2018). Importantly, our scRNA-seq data, as well as follow up experiments with *Ndufs4* knockdown macrophages *in vitro*, argue that mitochondrial respiratory chain collapse is concomitant with functional changes in macrophages that dampen antimycobacterial immune responses (**Fig. 5H-K**). These newly described defects in the ability of “exhausted” myeloid cells to express inflammatory mediators, present antigen, and constrain bacillary burden may help explain why Mtb can persist in mice long after they have mounted strong adaptive immune responses.

Precisely when this shift from high ETC gene expression to low ETC gene expression occurs and what triggers it remain unclear. Our *in vitro* experiments suggest that prolonged exposure to high levels of IL-1β and/or TLR2 stimulation are sufficient to reduce expression of ETC-associated genes (**Fig. 4E-H**). If a similar phenomenon occurs *in vivo*, it follows that levels of one or more inflammatory mediators cross this threshold sometime after day 21, triggering downregulation of ETC and glycolysis genes and subsequent bioenergetic collapse. Seminal studies have established that IL-1α/β protein levels peak in the lungs on Mtb infected mice at 4 weeks post infection (Mayer-Barber et al., 2011). Performing scRNA-seq analysis at infection timepoints closer to this tipping point would likely provide more resolution into how and when this myeloid cell metabolic reprogramming proceeds *in vivo*.

Although altered RNA abundance via scRNA-seq does not necessarily correlate with functional/phenotypic changes, our *in vitro* experiments clearly demonstrate that Mtb infection induces loss of mitochondrial ETC gene expression at the transcript and protein level (**Fig. 4A-C**) that result in changes to OXPHOS output and cellular ATP levels (**Fig. 4I-O**). These results strongly suggest that ETC low myeloid cells are defective in mitochondrial respiration *in vivo*. Further supporting this claim, SCENITH experiments conducted by Dkhar et al. suggest that AMs isolated from Mtb-infected mice are defective in their capacity to utilize mitochondrial OXPHOS to generate ATP compared to uninfected controls (Dkhar et al., 2026). Likewise, the Steyn lab recently reported that cellular bioenergetics are severely diminished in circulating monocytes from human pulmonary tuberculosis patients compared to healthy controls (Cumming et al., 2024).

A similar type of myeloid exhaustion has also been detected and characterized in other human disease states. Myeloid exhaustion was recently described in a mouse model of Parkinson’s disease, where mutations in LRRK2, a protein our labs have repeated linked to mitochondrial homeostasis, cause defects in myeloid antigen presentation, lysosomal function, and pathogen uptake (Wallings et al., 2024). Likewise, loss of ETC integrity and mitochondrial bioenergetic defects in conventional DC1s have been linked to tumor progression (You et al., 2026). Specifically, as tumors progress, cDC1s in the tumor microenvironment exhibit a decline in mitochondrial membrane potential and volume. It is tempting to speculate that the same inflammatory signals driving respiratory chain collapse in Mtb-infected mice promote parallel changes in tumor-associated DCs. Excitingly, introduction of cDC1s with high mitochondrial fitness was shown to promote T cell responses and tumor control, compared to cDC1s with depolarized mitochondria, suggesting that replenishing exhausted myeloid cells may also promote protective responses against Mtb.

Although the lower ETC transcriptional signatures observed in IGRA+ TB household contacts support the potential relevance of our mouse findings to human disease, it is impossible to conclude from these data alone how ETC low myeloid cells drive Mtb disease progression in humans. In our preferred model, household contacts that become IGRA+ possess intrinsically ETC-low myeloid cells, due to genetic, dietary, or other host factors. This ETC low signature renders them less capable of controlling Mtb following exposure, which results in T cell activation and seroconversion. Alternatively, IGRA+ individuals may be actively mounting an antimycobacterial response, such that these ETC-low monocytes represent a state analogous to that observed in our day 77 mice. Our data do not establish whether the ETC-low phenotype precedes or follows Mtb, nor do they definitively implicate ETC low monocytes in progression to active disease or protection from it. They do, however, identify mitochondrial respiratory chain remodeling as an important feature of the human myeloid response to tuberculosis and highlight the need for prospective studies that track individuals before exposure and throughout the course of infection.

## ACKNOWLEGEMENTS

We thank the members of the Watson and Patrick labs for their critical review and feedback in the preparation of this manuscript. We also thank Dr. Timothy Sterling, Dr. Bruno Andrade and other members of the RePORT-Brazil consortium for granting us access to scRNA-seq data from Penã Avila et al., 2025. Imaging was done with the help of the Vanderbilt Cell Imaging Shared Resource (supported by NIH grants CA68485, DK20593, DK58404, DK59637 and EY08126). Flow Cytometry experiments were performed in the VMC Flow Cytometry Shared Resource (supported by the Vanderbilt Ingram Cancer Center (P30 CA68485) and the Vanderbilt Digestive Disease Research Center (DK058404)). This work was supported in part by R01AI155621 to ROW and KLP, and R01AI179037 to ROW and KLP. Trainees were supported by F31AI197727 to ELM, F31AI176652 to AKC, F31AI176795 to CJM, F31CA298557 to MHS, F31AI197784 to KSA, 5T32GM137793 to JBH, and T32GM135748 to AKC.

## METHODS

### Experimental Models and Subject Details

#### Mouse husbandry and strains

C57BL6/J (Jackson laboratories stock # 000664) were purchased from Jackson Laboratories (Bar Harbor, ME). All mice used in experiments were compared to age- and sex-matched controls. Littermate controls were used in all experiments and fed a standard 4% chow diet. For *ex vivo* BMDM experiments, male mice 8-12 weeks old were used. For *in vivo* experiments, mice were used at 10-12 weeks. All animals were housed, bred, and studied at Texas A&M Health Science Center and Vanderbilt University Medical Center under approved Institutional Care and Use Committee guidelines.

#### M. tuberculosis

The Erdman strain (Erdman WT and Erdman mCherry) was used for *M. tuberculosis* infections. Low passage lab stocks were thawed for each experiment to ensure virulence was preserved. *M. tuberculosis* was cultured in roller bottles at 37°C in Middlebrook 7H9 broth (BD Biosciences) supplemented with 10% OADC (BD Biosciences), 0.5% glycerol (Fisher), and 0.1% Tween-80 (Fisher) or on 7H10 plates. All work with *M. tuberculosis* was performed under Biosafety level 3 containment using procedures approved by the Texas A&M and Vanderbilt University Institutional Biosafety Committee.

For *M. tuberculosis* infections *ex vivo*, low passage Mtb was prepared by growing it to log phase (OD_600_ 0.6–0.8). Bacterial cultures were spun at 58 rcf for 5 min to remove large clumps. The bacteria were then pelleted at 2,103 rcf for 5 min and washed with 1× PBS. The wash step was repeated twice. The resuspended bacterial cultures were sonicated at 70% amplitude for 10 s and repeated three times (Branson Ultrasonics Corp.) followed by a low-speed spin (58 rcf) to remove remaining clumps. The bacteria were diluted in DMEM (Hyclone) + 10% horse serum (Gibco) for *ex vivo* infections or 1× PBS for *in vivo* infections. For *ex vivo* infections, plates containing cells and bacteria were spun at 234 rcf for 10 min to synchronize infection. Fresh BMDM media was then added to the cells and replenished every 24h.

All Mtb infections were performed using procedures approved by Texas A&M University Institutional Care and Use Committee. The Mtb inoculum was prepared as described above. Age- and sex-matched mice were infected via inhalation exposure using a Madison chamber (Glas-Col) calibrated to introduce 100-200 CFUs per mouse. For each infection, approximately 3 mice were euthanized immediately, and their lungs were homogenized and plated to verify an accurate inoculum. Infected mice were housed under BSL3 containment and monitored daily by lab members and veterinary staff. At the indicated time points, mice were euthanized, and tissue samples were collected. Organs were further processed as described below. For CFU enumeration, organs were homogenized in 5 ml PBS + 0.1% Tween-80, and serial dilutions were plated on 7H10 plates. Colonies were counted after plates were incubated at 37°C for 3 weeks.

### Primary and immortalized cell culture

Bone marrow derived macrophages were differentiated from bone marrow isolated from mouse femurs with DMEM supplemented with 1mM sodium pyruvate as in (Weindel et al., 2022).

Immortalized BMDMs (iBMDMs) were generated with the Cre-J2 virus as described in (De Nardo et al., 2018). iBMDMs were cultured in DMEM, 10% FBS, and 2% HEPES (Cytiva) and grown at 37°C 5% CO2.

### Isolation of single cell suspensions from uninfected mouse lungs

Single-cell lung suspensions were isolated from uninfected mice with a lung dissociation kit (Miltenyi Biotec) and gentleMACS tissue dissociator (Miltenyi Biotec) per manufacturer’s protocol. CD45^+^ cell populations were enriched in pooled cell suspensions using CD45 magnetic microbeads (Miltenyi Biotec) and a Miltenyi magnetic separator per manufacturer’s protocol. After magnetic enrichment, CD45^+^ cell populations were processed for single cell 3’ RNA sequencing utilizing the 10X Chromium system (10x Genomics). Sequencing was performed at the VUMC Vanderbilt Technologies for Advanced Genomics (VANTAGE) core on the Illumina NovaSeq XP sequencer.

### Isolation of single cell suspensions from Mtb infected mouse lungs

Mtb-infected mouse left lung lobe was isolated and washed twice in 1x PBS. The lung was then minced and digested in digestion buffer (70 µg/mL Liberase [Roche], 50 µg/mL DNase I [Worthington Biochemical] in RPMI 1640 [HyClone], and 55 µM 2-mercaptoethanol [Thermo Fisher Scientific]) for 30 min at 37°C and 5% CO_2_. Single suspensions were achieved by consecutively filtering homogenates through 70 µm and 40 µm cell strainers. CD45^+^ cell populations were enriched in pooled cell suspensions using CD45 magnetic microbeads (Miltenyi Biotec) and a Miltenyi magnetic separator per manufacturer’s protocol. After magnetic enrichment, CD45^+^ cell populations were methanol fixed per manufacturer’s protocol (10x Genomics) and processed for single cell 3’ RNA sequencing utilizing the 10X Chromium system (10x Genomics). Sequencing was performed at the Texas A&M University Molecular Genomics Core on the Illumina NextSeq 2000 sequencer.

### Single cell RNA sequencing

Sample demultiplexing, barcode processing and counting was performed using the 10x Genomics Cloud Analysis platform. Cell Ranger Count v9.0.1 was used to align samples to the Mouse (GRCm39) 2024-A reference genome, filtering and quantifying reads. The filtered feature-barcode matrices were used for downstream analysis.

### Single cell RNA sequencing processing and quality control

Single cell RNA sequencing data were analyzed in R 4.5.2 using Seurat 5.4.0. Filtered count matrices were imported into Seurat, retaining genes detected in at least 3 cells and cells with at least 200 detected features. Doublets were identified independently within each library using scDblFinder 1.24.10 and cells classified as singlets were retained for downstream analysis. Libraries were subsequently merged, and cells with ≤200 or ≥10,000 detected genes or ≥15% mitochondrial transcripts were excluded. Data were normalized using SCTransform with mitochondrial transcript percentage included as a regression variable. Principal component analysis was performed on the SCTransform normalized data, followed by Harmony integration. UMAP visualization was generated using the first 50 Harmony dimensions with a minimum distance of 0.3, and nearest-neighbor graphs were constructed from the Harmony embedding using the FindNeighbors function and graph based clustering was constructed using on the Harmony reduction using FindClusters, both functions provided by Seurat (Butler et al., 2018).

### Cell clustering and annotation

For the initial merged atlas, clustering using the provided FindClusters function was evaluated across resolutions from 0 to 2 in increments of 0.2, with a resolution of 0.6 used for broad cell type annotation. Cluster enriched genes were identified on the SCT assay using FindAllMarkers with a minimum log2 fold change threshold of 0.25 and expression in at least 25% of cells. Populations were annotated manually using canonical linages and cell type markers. Myeloid cells were subsequently subset and reclustered, and final myeloid identities used for downstream analyses were defined at resolution 0.9. Erythroid populations, stressed cell clusters and contaminating T cell populations were excluded from the final myeloid dataset.

### Differential Expression analysis and pathway enrichment

Differential expression between experimental conditions were performed using FindMarkers on the SCT assay following PrepSCTFindMarkers function. Average log2 fold was used as the gene level effect size and as the ranking statistic for pre-ranked gene set enrichment analysis (GSEA).

GSEA was performed using fgeaMultilevel in fgsea 13.36.0. Gene sets were obtained from MsigDB hallmark, Reactome and Gene ontology Biological Process collections. Genes were ranked in decreasing order of average log2FC. Gene sets containing 15-1000 genes were evaluated with nPermSimple = 10000 and pathway significances was assessed using Benjamini-Hochberg adjusted p values, with adjusted p values <0.05 considered significant. Normalized enrichments score (NES) was used to describe the direction and magnitude of pathway enrichment.

### OXPHOS gene expression and module score analyses

For the differential expression heatmap, log2 fold changes were standardized separately within each comparison where each gene fold change is relative to mean gene fold change for each comparison. These values therefore represent the relative magnitude and direction of differential expression within each comparison, rather than absolute or scaled gene expression for Day 21 versus uninfected, Day 77 versus uninfected, and Day 77 versus Day 21.

For expression based heatmaps, expression of genes was summarized across infection conditions using SeuratExtend::CalcStats, with samples as the grouping variable. Values represent gene-wise standardized expression across conditions, such that each gene is scaled relative to its own expression across samples.

Representative OXPHOS module score was calculated for each myeloid cell using Seurat’s AddModuleScore with *Ndufs4*, *Ndufa13*, *Sdhb*, *Sdhc*, *Uqcrb*, *Uqcrc2*, *Cox7a2*, *Cox4i1*, *Atp5d*, and *Atp5e*, representing OXPHOS complexes I–V. Representative genes were also visualized across myeloid populations and infection conditions using scaled gene expression.

Cells were additionally classified as low, intermediate, or high OXPHOS using 0-25^th^, 25^th^-75^th^, and 75^th^ -100^th^ percentiles respectively of the OXPHOS score distribution.

### Identification of transcriptional programs associated with OXPHOS

To identify transcriptional programs associated with OXPHOS state independently within distinct myeloid populations, Spearman’s rank correlation was calculated between SCT-normalized expression of each gene and the single-cell OXPHOS module score. Correlations were calculated separately within each annotated myeloid population using all cells and independently for Day 21 and Day 77 cells. Genes were ranked by Spearman’s ρ and subjected to pre-ranked GSEA using Hallmark, Reactome, and GO Biological Process gene sets. Positive NES values identify transcriptional programs positively correlated with OXPHOS scores, whereas negative NES values identify programs associated negatively correlated with OXPHOS scores.

### Consensus Pathway signatures and OXPHOS-state analysis

For selected OXPHOS associated pathways, consensus leading-edge signatures were generated from the Day 77 correlation-based GSEA. Pathways significant in at least two myeloid populations were considered. For each pathway, genes were retained when they occurred in the GSEA leading edge of at least 50% of the cell types in which that pathway was significant, and signatures containing fewer than 5 consensus genes were excluded. Consensus pathway module scores were then calculated at the single cell level using AddModuleScore

Two related OXPHOS decile analyses were performed. To examine redistribution of cells across OXPHOS states, Day 21 and Day 77 cells were jointly divided into OXPHOS score deciles within each myeloid population. For each cell type and time point, the proportion of cells occupying each decile was calculated, and these cell-type specific proportions were averaged across populations to obtain mean decile occupancy. This approach gives each annotated myeloid population equal contribution to the aggregate distribution rather than weighting the result by population abundance.

For analyses relating functional pathway scores to OXPHOS state, deciles were instead defined independently within each myeloid population and time point, ensuring representation across the complete within population OXPHOS distribution at both Day 21 and Day 77. Within each cell type, day, pathway, and decile, median OXPHOS and pathway module scores were calculated. These cell-type specific medians were then averaged across myeloid populations. Accordingly, plotted values represent cell-type averaged summaries, rather than pooled cell or abundance weighted estimates.

Spearman correlations between the cell-type averaged OXPHOS and pathway scores were calculated separately at Day 21 and Day 77. Differences between time points were further evaluated using linear models of the form Pathway Module score ∼ OXPHOS score × Day

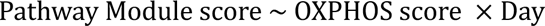

from which day and Day by OXPHOS interaction terms were evaluated.

### Effect Size Analysis

The magnitude and direction of temporal changes in pathway module score were summarized within each myeloid population using Cohen’s d. This was used to compare the magnitude of day 21 to day 77 changes across myeloid populations on a common scale. This was useful because different populations have different baseline pathway scores and levels of cell-to-cell variability (Lakens, 2013). Effect sizes were calculated as

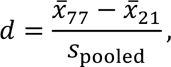

where

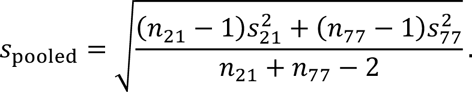

Positive values therefore indicate higher pathway module scores at Day 77 relative to Day 21, whereas negative values indicate lower scores. Cohen’s *d* was used as a descriptive standardized measure of the magnitude differences in single-cell score distributions and was not interpreted as a mouse-level effect estimate.

### Seahorse metabolic analysis

BMDMs were plated in 24-well Seahorse cell culture microplates at 2×10^5^ cells/well. The following day, BMDMs were infected as described above at indicated MOIs (multiplicity of infection) and timepoints. The Seahorse XF mito stress test kit and cartridge were prepared per Agilent’s protocols with exception of the first injection with glucose as described in (Van den Bossche et al., 2015; Weindel et al., 2022). Normalization was based on bicinchoninic acid assay (BCA) absorbance values (Thermo Fisher Scientific).

### Protein analysis by immunoblot

BMDMs were seeded in 12-well plates at 5×10^5^ cells/well. At indicated timepoints, cells were lysed in 1% SDS lysis buffer. Protein quantification was measured using BCA. 30 ug of protein was separated by SDS-PAGE and transferred to 0.45 µM PVDF-FL membranes (Millipore-Sigma). Membranes were blocked for 1h at room temperature in 5% BSA in 1X TBS. Blot were incubated overnight at 4°C with the following antibodies: ACTIN (Cell Signaling Technology (CST) 4970, 1:1000), NDUFS4 (Proteintech 15849-1-AP, 1:1000), NDUFB8 (CST 73951, 1:1000), OXPHOS (Abcam ab110413, 1:1000), and COXIV (CST 11967, 1:1000). All washes were done with 1X TBST (0.1% Tween-20). Membranes were incubated with appropriate LiCOR fluorescent secondary antibodies in 5% BSA/TBS for 2h at RT prior to imaging on a LiCOR Odyssey Fc Imager.

### Cell stimulation/agonist treatments

Murine BMDMs were stimulated using 1 ng/mL IL-1β (Abcam) or 20 ng/mL Pam3CSK4 (Invivogen) for one, three, or five days respectively in media containing DMEM, 20% heat-inactivated FBS, and 10% MCSF. Media containing the agonist or cytokine was replaced every 24 hours. RNA was extracted in TRIzol and purified with Direct-zol RNA Miniprep kits (Zymo Research) including on-column DNase digestion. Following RNA extraction, cDNA synthesis was performed using iScript cDNA Synthesis Kit (Bio-Rad). qPCR was performed with PowerUp SYBR Green Master Mix (Thermo Fisher) on a QuantStudio Flex6 instrument.

### iBMDM shRNA KDs

For iBMDMs stably expressing scramble knockdown and *Ndufs4* knockdown, Lenti-X cells were transfected with a pSICO-R scramble non-targeting shRNA construct or pSICO *Ndufs4* shRNA constructs using Polyjet (SignaGen Laboratories). Virus was collected 24 and 48h post transfection and used to transduce iBMDMs. After 48h, media was supplemented with Hygromycin (Invitrogen) to select for cells containing the shRNA plasmid.

### RNA Sequencing and Bioinformatics Analysis Differential Expression Analysis

RNA was extracted from infected and uninfected BMDM samples in triplicate or quadruplicate using the Zymo Research Direct-zol RNA Kit (R2052). RNA sequencing was performed by Plasmidsaurus using 3’ end counting using an Illumina NovaSeq X Plus with custom analysis and annotation. In-house scripts were used to analyze the counts data obtained from Plasmidsaurus. Additional differential expression analysis was performed in R (v4.5.2) using the DESeq2 (v1.50.2) package. Raw, un-normalized transcript count data and sample metadata were imported, and sample identifies were cross-referenced to ensure exact matrix alignment. Low-abundance transcripts were filtered out by removing any gene failing to achieve a total sum of 10 or more raw reads across all samples. Raw p-values were adjusted using the Benjamini-Hochberg False Discovery Rate (FDR) procedure to generate the adjusted p-value (Love et al., 2014).

### Gene Set Enrichment Analysis

Permutation testing was executed against three distinct databases sourced from the Molecular Signatures Database (MSigDB) [v25.1.1] *Mus musculus*. The Gene Ontology Biological Processes (GO:BP) category C5, Hallmark Pathways, and Canonical Reactome Pathways. To eliminate overly broad or hyper-specific functional annotations, pathway enrichment evaluations were strictly constrained to gene sets containing a minimum of 15 and a maximum of 500 genes. Resulting pathways were sorted and prioritized by their Benjamini-Hochberg adjusted p-values (padj) of *p-adj* < 0.05 (Martin Morgan et al., 2026).

### Pathway-Specific Expression Heatmaps

To visualize transcriptomic variation within prioritized pathways across experimental conditions, normalized gene expression metrics were extracted for specific target gene sets. Sample expression data were standardized for downstream analysis using a variance-stabilizing transformation (VST) from DESeq2. The VST values for the targeted genes were subsequently transformed into cross-sample z-scores to standardize baseline variation across features. Heatmaps were generated using the ComplexHeatmap R package (v2.26.1) with hierarchical clustering on genes (Gu, 2022; Gu et al., 2016).

### Volcano plots

To visualize the global transcriptomic profile and isolate individual candidate genes, volcano plots were generated using ggplot2 (v4.0.2) (Wickham, 2016) while highlighting points with a padj < 0.05 and the absolute value of the log_2_Fold Change greater than 1. Statistically significant candidates were labeled using ggrepel (v0.9.6)(Slowikowski, 2026).

### Immunofluorescence microscopy

BMDM cells were seeded at 2.5×10^5^ cells/well on glass coverslips in 24-well dishes and infected with WT Mtb at MOI=1 for 1, 3, and 5 days. For a positive control to induce MHC Class II expression, cells were stimulated with 50 ng/ml IFNγ for 24h. Cells were fixed in 4% PFA for 15 min at RT and then washed three times with PBS. Coverslips were incubated in the primary antibody diluted in PBS + 5% non-fat milk +0.1%Triton-X (PBS-MT) for 3 hrs. Primary antibody used in this study was Anti-Mycobacterium tuberculosis (Abcam, ab905,1:100). Cells were then washed three times in PBS and incubated in secondary antibodies (goat anti-rabbit Alexa Fluor 488; Invitrogen, A11008, 1:500) PBS-MT for 1 hr. Coverslips were washed twice with 1X PBS and twice with deionized water and mounted on glass slides using Fluoromount-G™ Mounting Medium, with DAPI (Invitrogen 00-4959-52). Z-stack images were obtained using a Zeiss LSM710 inverted confocal microscope equipped with 40X water immersion objective for quantitated images and a 60X oil immersion objective for representative images with DIC channel for both. Quantifications were performed using Fiji ImageJ. Representative images were opened as separate channels and z-stacks were converted to maximum intensity projections.

To measure MHCII expression, iBMDM SCR and Ndufs4 KD cells were seeded at 2.5×10^5^ cells/well on glass coverslips in 24-well dishes. Cells were fixed in 4% PFA for 10 min at RT and then washed three times with PBS. Coverslips were incubated in the primary antibody diluted in PBS + 5% non-fat milk +0.1%Triton-X (PBS-MT) for 3 hrs. Conjugated primary antibody used in this study was APC anti-mouse I-A/I-E, MHC class II, M5/114.15.2 (Biolegend, 107614, 1:100). Coverslips were washed twice with 1X PBS and twice with deionized water and mounted on glass slides using Fluoromount-G™ Mounting Medium, with DAPI (Invitrogen 00-4959-52). Z-stack images were obtained using an Zeiss LSM710 inverted confocal microscope equipped with 40X water immersion objective for representative images and a 60X oil immersion objective for quantitative images with DIC channel for both. Quantifications were performed using Fiji ImageJ. Representative images were opened as separate channels and z-stacks were converted to maximum intensity projections.

To follow Mtb infection, iBMDM SCR and Ndufs4 KD cells were seeded at 250,000 cells/well on glass coverslips in 24-well dishes and infected with Erdman Mtb at an MOI=1 for 5 days. Cells were fixed in 4% PFA for 10 min at RT and then washed three times with PBS. Coverslips were permeabilized in PBS + 5% non-fat milk +0.1%Triton-X (PBS-MT) for 10 minutes. Coverslips were washed twice with 1X PBS and twice with deionized water and mounted on glass slides using Fluoromount-G™ Mounting Medium, with DAPI (Invitrogen 00-4959-52). Z-stack images were obtained using a Zeiss LSM710 inverted confocal microscope equipped with 40X water immersion objective for quantitated images and a 60X oil immersion objective for representative images with DIC channel for both. Quantifications were performed using Fiji ImageJ. Representative images were opened as separate channels and z-stacks were converted to maximum intensity projections.

### Software and statistical analysis

Analyses were performed in R 4.5.2. Major packages included Seurat 5.4.0, SeuratObject 5.3.0, Harmony 2.0.2, scDblFinder 1.24.10, SeuratExtend 1.2.8, fgsea 1.36.0, msigdbr 25.1.1, ComplexHeatmap 2.26.0, rstatix 0.7.3, ggplot2 4.0.1, dplyr 1.1.4, and broom 1.0.12.

