## Supplemental Figures S1-S6 for "Loss of Mitochondrial Respiratory Capacity Reshapes Myeloid Cell Function during *Mycobacterium tuberculosis* infection"

#### SUPPLEMENTAL FIGURE LEGENDS

**Figure S1**

(A) UMAP visualization of aggregate data representing the total immune landscape of Mtb infected mouse lung samples (uninfected, day 21 and day 77).

(B) Dot plot showing representative marker genes for each cell population. Dot size indicates percentage of cells expressing each gene and color intensity represents scaled gene expression (Z-score). Colors denote broad myeloid categories.

(C) UMAP visualization of myeloid immune landscape of Mtb infected mouse lung split by sample (uninfected, day 21 and day 77).

(D) Bar plot showing relative abundance of cells across experimental conditions. Bars show percentage of cells contributed by each sample within each cell population.

(E) Dot plot showing Log2 fold change between in cell number for each population between day 77 and day 21. Positive values (red) indicate increased representation at day 77 whereas negative values (blue) indicate decreased representation

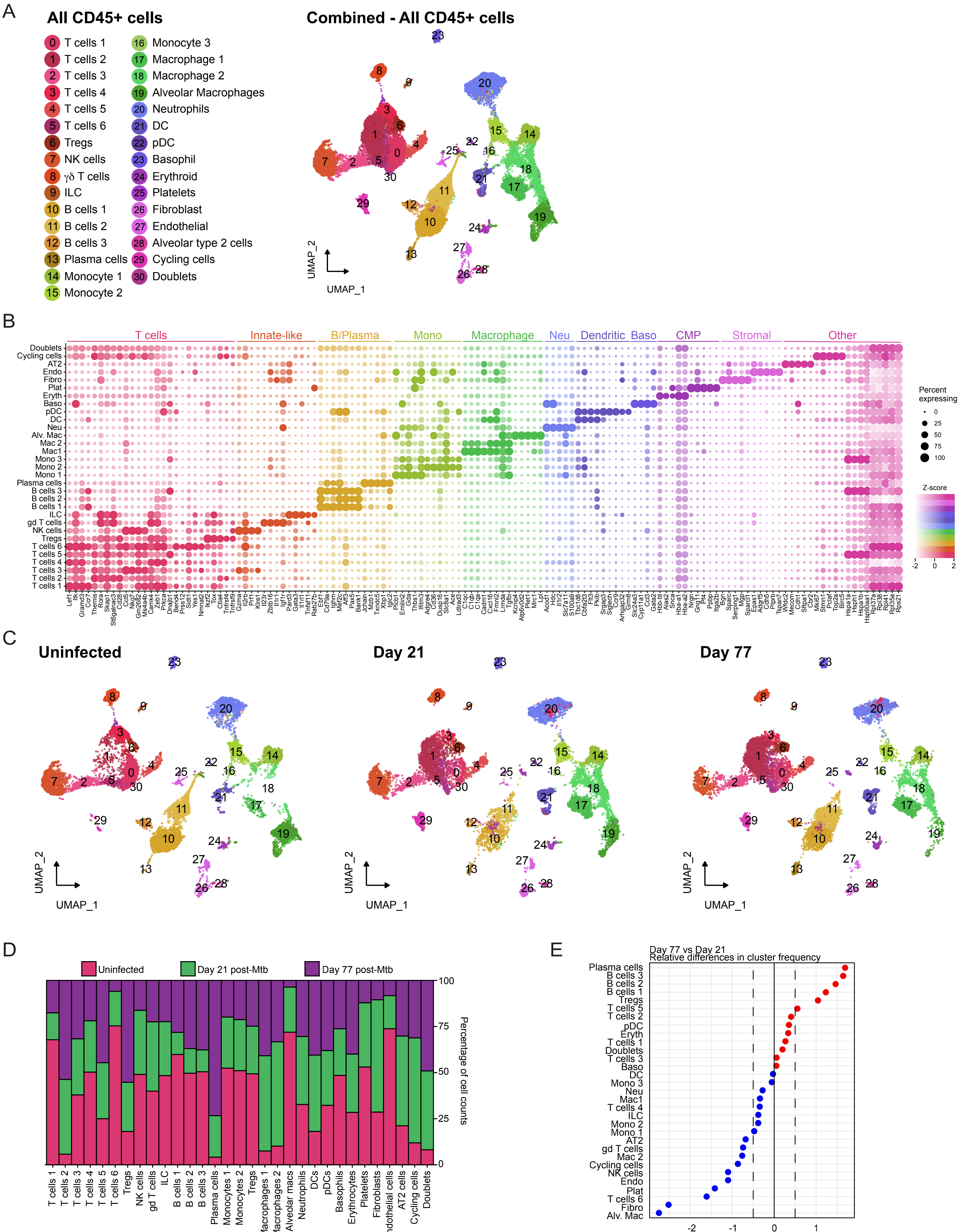

A
All CD45+ cells
0 T cells 1
1 T cells 2
2 T cells 3
3 T cells 4
4 T cells 5
5 T cells 6
6 Tregs
7 NK cells
8 γδ T cells
9 ILC
10 B cells 1
11 B cells 2
12 B cells 3
13 Plasma cells
14 Monocyte 1
15 Monocyte 2
16 Monocyte 3
17 Macrophage 1
18 Macrophage 2
19 Alveolar Macrophages
20 Neutrophils
21 DC
22 pDC
23 Basophil
24 Erythroid
25 Platelets
26 Fibroblast
27 Endothelial
28 Alveolar type 2 cells
29 Cycling cells
30 Doublets
Combined - All CD45+ cells
UMAP_2
UMAP_1
B
T cells
Innate-like
B/Plasma
Mono
Macrophage
Neu
Dendritic
Baso
CMP
Stromal
Other
Doublets
Cycling cells
AT2
Endo
Fibro
Plat
Eryth
Baso
pDC
DC
Neu
Alv. Mac
Mac 2
Mac1
Mono 3
Mono 2
Mono 1
Plasma cells
B cells 3
B cells 2
B cells 1
ILC
gd T cells
NK cells
Tregs
T cells 6
T cells 5
T cells 4
T cells 3
T cells 2
T cells 1
Percent expressing
0
25
50
75
100
Z-score
-2
0
2
C
Uninfected
Day 21
Day 77
UMAP_2
UMAP_1
D
Uninfected
Day 21 post-Mtb
Day 77 post-Mtb
Percentage of cell counts
100
75
50
25
0
T cells 1
T cells 2
T cells 3
T cells 4
T cells 5
T cells 6
Tregs
NK cells
gd T cells
ILC
B cells 1
B cells 2
B cells 3
Plasma cells
Monocytes 1
Monocytes 2
Tregs
Macrophages 1
Macrophages 2
Alveolar macs
Neutrophils
DCs
pDCs
Basophils
Erythrocytes
Platelets
Fibroblasts
Endothelial cells
AT2 cells
Cycling cells
Doublets
E
Day 77 vs Day 21
Relative differences in cluster frequency
Plasma cells
B cells 3
B cells 2
B cells 1
Tregs
T cells 5
T cells 2
pDC
Eryth
T cells 1
Doublets
T cells 3
Baso
DC
Mono 3
Neu
Mac1
T cells 4
ILC
Mono 2
Mono 1
AT2
gd T cells
Mac 2
Cycling cells
NK cells
Endo
Plat
T cells 6
Fibro
Alv. Mac
-2
-1
0
1
log2FC


Figure S1

**Figure S2**

(A-B) Pathway analysis for aggregate myeloid day 21 vs uninfected and day 77 vs uninfected respectively. Dot size indicates gene set size and color indicates positive (red) or negative (blue) enrichment for indicated days post infection.

(C) Heatmap of genes encoding OXPHOS complexes I-V, grouped by respiratory complex. Values represent row-wise scaled expression (z-score) across samples.

(D) trajectory plot showing median OXPHOS score trajectories for individual myeloid populations, lines connect the median score for each population at uninfected, day 21 and day 77.

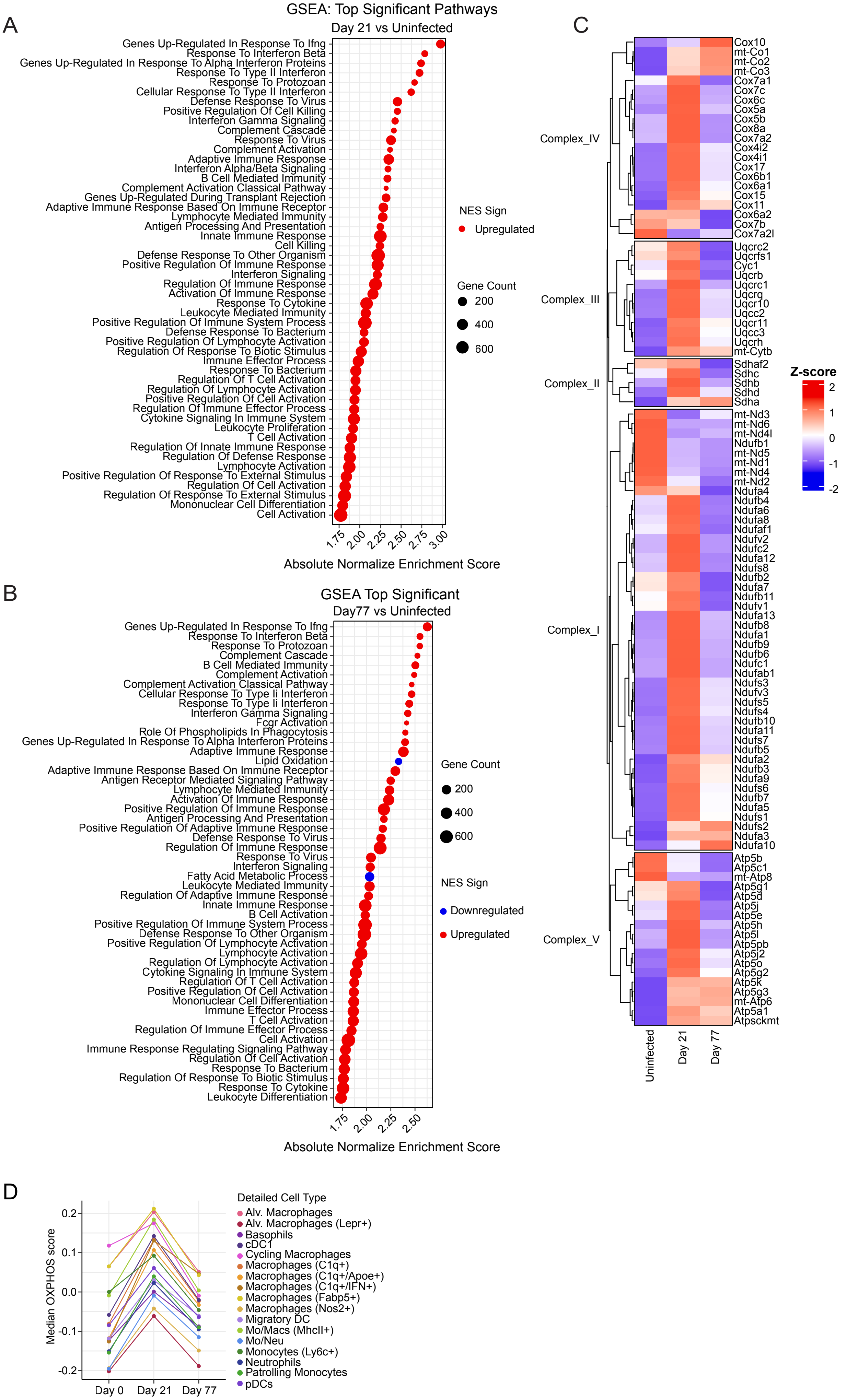
A
GSEA: Top Significant Pathways
Day 21 vs Uninfected
Genes Up-Regulated In Response To Ifng
Response To Interferon Beta
Genes Up-Regulated In Response To Alpha Interferon Proteins
Response To Type II Interferon
Response To Protozoan
Cellular Response To Type II Interferon
Defense Response To Virus
Positive Regulation Of Cell Killing
Interferon Gamma Signaling
Complement Cascade
Response To Virus
Complement Activation
Adaptive Immune Response
Interferon Alpha/Beta Signaling
B Cell Mediated Immunity
Complement Activation Classical Pathway
Genes Up-Regulated During Transplant Rejection
Adaptive Immune Response Based On Immune Receptor
Lymphocyte Mediated Immunity
Antigen Processing And Presentation
Innate Immune Response
Cell Killing
Defense Response To Other Organism
Positive Regulation Of Immune Response
Interferon Signaling
Regulation Of Immune Response
Activation Of Immune Response
Response To Cytokine
Leukocyte Mediated Immunity
Positive Regulation Of Immune System Process
Defense Response To Bacterium
Positive Regulation Of Lymphocyte Activation
Regulation Of Response To Biotic Stimulus
Immune Effector Process
Response To Bacterium
Regulation Of T Cell Activation
Regulation Of Lymphocyte Activation
Positive Regulation Of Cell Activation
Regulation Of Immune Effector Process
Cytokine Signaling In Immune System
Leukocyte Proliferation
T Cell Activation
Regulation Of Innate Immune Response
Regulation Of Defense Response
Lymphocyte Activation
Positive Regulation Of Response To External Stimulus
Regulation Of Cell Activation
Regulation Of Response To External Stimulus
Mononuclear Cell Differentiation
Cell Activation
1.75 2.00 2.25 2.50 2.75 3.00
Absolute Normalize Enrichment Score
NES Sign
Upregulated
Gene Count
200
400
600
B
GSEA Top Significant
Day77 vs Uninfected
Genes Up-Regulated In Response To Ifng
Response To Interferon Beta
Response To Protozoan
Complement Cascade
B Cell Mediated Immunity
Complement Activation
Complement Activation Classical Pathway
Cellular Response To Type Ii Interferon
Response To Type Ii Interferon
Interferon Gamma Signaling
Fcgr Activation
Role Of Phospholipids In Phagocytosis
Genes Up-Regulated In Response To Alpha Interferon Proteins
Adaptive Immune Response
Lipid Oxidation
Adaptive Immune Response Based On Immune Receptor
Antigen Receptor Mediated Signaling Pathway
Lymphocyte Mediated Immunity
Activation Of Immune Response
Positive Regulation Of Immune Response
Antigen Processing And Presentation
Positive Regulation Of Adaptive Immune Response
Defense Response To Virus
Regulation Of Immune Response
Response To Virus
Interferon Signaling
Fatty Acid Metabolic Process
Leukocyte Mediated Immunity
Regulation Of Adaptive Immune Response
Innate Immune Response
B Cell Activation
Positive Regulation Of Immune System Process
Defense Response To Other Organism
Positive Regulation Of Lymphocyte Activation
Lymphocyte Activation
Regulation Of Lymphocyte Activation
Cytokine Signaling In Immune System
Regulation Of T Cell Activation
Positive Regulation Of Cell Activation
Mononuclear Cell Differentiation
Immune Effector Process
T Cell Activation
Regulation Of Immune Effector Process
Cell Activation
Immune Response Regulating Signaling Pathway
Regulation Of Cell Activation
Response To Bacterium
Regulation Of Response To Biotic Stimulus
Response To Cytokine
Leukocyte Differentiation
1.75 2.00 2.25 2.50
Absolute Normalize Enrichment Score
Gene Count
200
400
600
NES Sign
Downregulated
Upregulated
C
Complex_IV
Cox10
mt-Co1
mt-Co2
mt-Co3
Cox7a1
Cox7c
Cox6c
Cox5a
Cox5b
Cox8a
Cox7a2
Cox4i2
Cox4i1
Cox17
Cox6b1
Cox6a1
Cox15
Cox11
Cox6a2
Cox7b
Cox7a2l
Complex_III
Uqcrc2
Uqcrfs1
Cyc1
Uqcrb
Uqcrc1
Uqcrq
Uqcr10
Uqcc2
Uqcr11
Uqcc3
Uqcrh
mt-Cytb
Complex_II
Sdhaf2
Sdhc
Sdhb
Sdhd
Sdha
Complex_I
mt-Nd3
mt-Nd6
mt-Nd4l
Ndufb1
mt-Nd5
mt-Nd1
mt-Nd4
mt-Nd2
Ndufa4
Ndufb4
Ndufa6
Ndufa8
Ndufaf1
Ndufv2
Ndufc2
Ndufa12
Ndufs8
Ndufb2
Ndufa7
Ndufb11
Ndufv1
Ndufa13
Ndufb8
Ndufa1
Ndufb9
Ndufb6
Ndufc1
Ndufab1
Ndufs3
Ndufv3
Ndufs5
Ndufs4
Ndufb10
Ndufa11
Ndufs7
Ndufb5
Ndufa2
Ndufb3
Ndufa9
Ndufs6
Ndufb7
Ndufa5
Ndufs1
Ndufs2
Ndufa3
Ndufa10
Complex_V
Atp5b
Atp5c1
mt-Atp8
Atp5g1
Atp5d
Atp5j
Atp5e
Atp5h
Atp5l
Atp5pb
Atp5j2
Atp5o
Atp5g2
Atp5k
Atp5g3
mt-Atp6
Atp5a1
Atpsckmt
Uninfected
Day 21
Day 77
Z-score
2
1
0
-1
-2
D
Median OXPHOS score
0.2
0.1
0.0
-0.1
-0.2
Day 0
Day 21
Day 77
Detailed Cell Type
Alv. Macrophages
Alv. Macrophages (Lepr+)
Basophils
cDC1
Cycling Macrophages
Macrophages (C1q+)
Macrophages (C1q+/Apoe+)
Macrophages (C1q+/IFN+)
Macrophages (Fabp5+)
Macrophages (Nos2+)
Migratory DC
Mo/Macs (MhcII+)
Mo/Neu
Monocytes (Ly6c+)
Neutrophils
Patrolling Monocytes
pDCs

Figure S2

**Figure S3**

(A) Top 20 positive and negative pathway GSEA. Genes were ranked by their Spearman correlation with OXPHOS module score within individual myeloid populations. Pathways significantly enriched in at least two myeloid populations are summarized by mean normalized enrichment score (NES) and dot size represents the number of cell types the pathway was significant.

(B) Line plots show the relationship between OXPHOS state and OXPHOS module scores for indicated pathways at day 21 and 77. Values represent cell type weighted summaries. Spearman correlations were calculated separately for each day, and linear models test day and day by OXPHOS interactions. Accompanying the lines plots are lollipop plots show the standardized differences in pathway module scores between day 77 and day 21 across individual populations. Cohen’s d was calculated as day 77 relative to day 21; positive values (red) indicate higher module scores at day 77 while negative values (blue) indicate lower module scores.

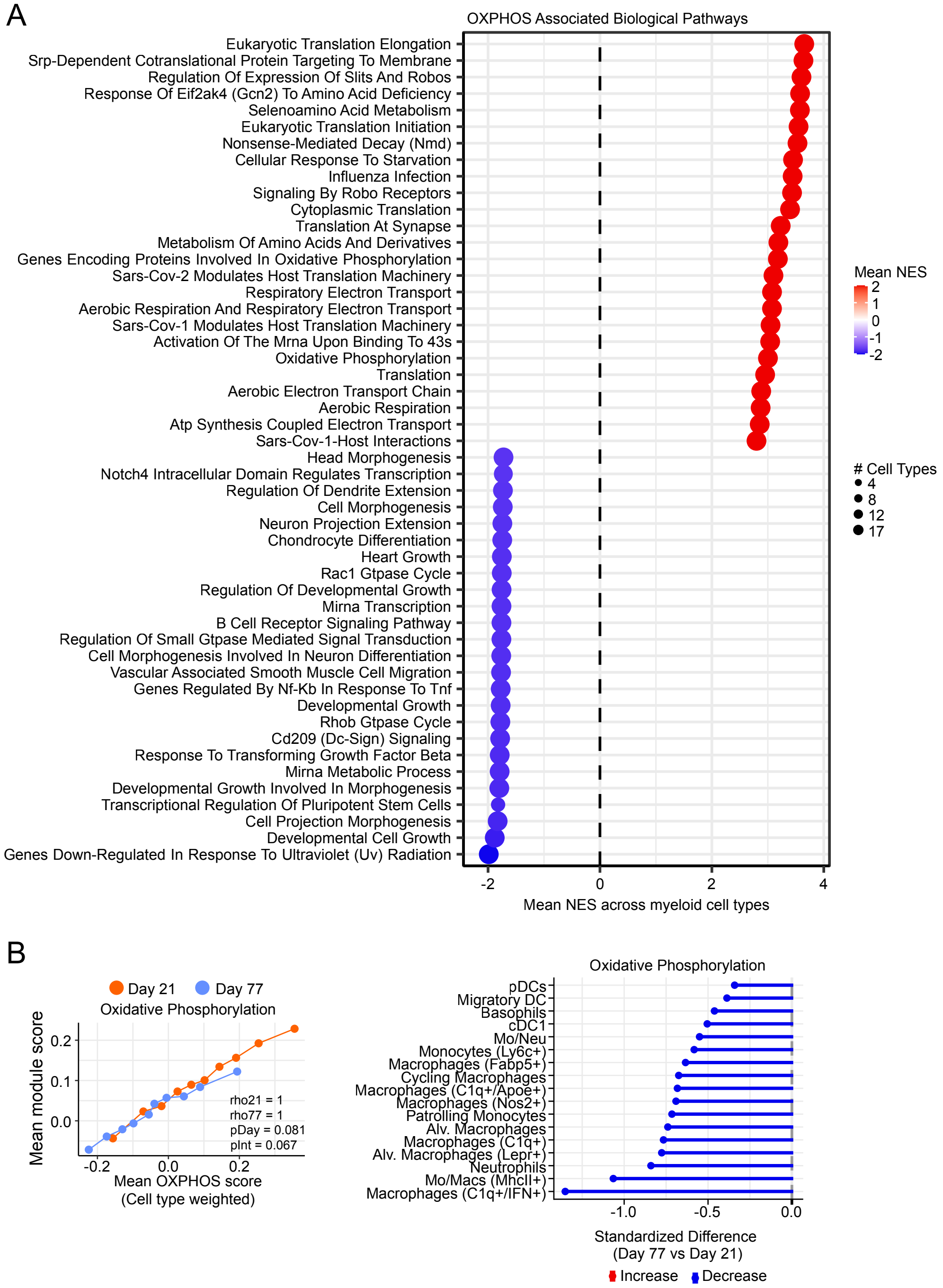
A
OXPHOS Associated Biological Pathways
Eukaryotic Translation Elongation
Srp-Dependent Cotranslational Protein Targeting To Membrane
Regulation Of Expression Of Slits And Robos
Response Of Eif2ak4 (Gcn2) To Amino Acid Deficiency
Selenoamino Acid Metabolism
Eukaryotic Translation Initiation
Nonsense-Mediated Decay (Nmd)
Cellular Response To Starvation
Influenza Infection
Signaling By Robo Receptors
Cytoplasmic Translation
Translation At Synapse
Metabolism Of Amino Acids And Derivatives
Genes Encoding Proteins Involved In Oxidative Phosphorylation
Sars-Cov-2 Modulates Host Translation Machinery
Respiratory Electron Transport
Aerobic Respiration And Respiratory Electron Transport
Sars-Cov-1 Modulates Host Translation Machinery
Activation Of The Mrna Upon Binding To 43s
Oxidative Phosphorylation
Translation
Aerobic Electron Transport Chain
Aerobic Respiration
Atp Synthesis Coupled Electron Transport
Sars-Cov-1-Host Interactions
Head Morphogenesis
Notch4 Intracellular Domain Regulates Transcription
Regulation Of Dendrite Extension
Cell Morphogenesis
Neuron Projection Extension
Chondrocyte Differentiation
Heart Growth
Rac1 Gtpase Cycle
Regulation Of Developmental Growth
Mirna Transcription
B Cell Receptor Signaling Pathway
Regulation Of Small Gtpase Mediated Signal Transduction
Cell Morphogenesis Involved In Neuron Differentiation
Vascular Associated Smooth Muscle Cell Migration
Genes Regulated By Nf-Kb In Response To Tnf
Developmental Growth
Rhob Gtpase Cycle
Cd209 (Dc-Sign) Signaling
Response To Transforming Growth Factor Beta
Mirna Metabolic Process
Developmental Growth Involved In Morphogenesis
Transcriptional Regulation Of Pluripotent Stem Cells
Cell Projection Morphogenesis
Developmental Cell Growth
Genes Down-Regulated In Response To Ultraviolet (Uv) Radiation
-2
0
2
4
Mean NES across myeloid cell types
Mean NES
2
1
0
-1
-2
### Cell Types
4
8
12
17
B
Day 21
Day 77
Oxidative Phosphorylation
Mean module score
0.2
0.1
0.0
-0.2
0.0
0.2
rho21 = 1
rho77 = 1
pDay = 0.081
pInt = 0.067
Mean OXPHOS score
(Cell type weighted)
Oxidative Phosphorylation
pDCs
Migratory DC
Basophils
cDC1
Mo/Neu
Monocytes (Ly6c+)
Macrophages (Fabp5+)
Cycling Macrophages
Macrophages (C1q+/Apoe+)
Macrophages (Nos2+)
Patrolling Monocytes
Alv. Macrophages
Macrophages (C1q+)
Alv. Macrophages (Lepr+)
Neutrophils
Mo/Macs (MhcII+)
Macrophages (C1q+/IFN+)
-1.0
-0.5
0.0
Standardized Difference
(Day 77 vs Day 21)
Increase
Decrease

**Figure S4**

A) Immunoblot analysis of Atp5a, Uqcrc2, mt-Co1, Sdhb in uninfected and Mtb-infected (MOI=1) BMDMs at 1, 3, and 5 days. Actin was used as a loading control.

B) Quantification of Sdhb protein levels normalized to Actin from A. n=3

C) As in B, but with Uqcrc2 protein levels normalized to Actin. n=3

D) As in B, but with Atp5a protein levels normalized to Actin. n=3

E) Basal respiration, maximal respiration, and spare respiratory capacity, measured in uninfected and Mtb-infected (MOI=1) BMDMs at 1, 3, and 5 days.

F) Glycolysis, maximal glycolytic capacity, and glycolytic reserve measured in uninfected and Mtb-infected (MOI=1) BMDMs at 1, 3, and 5 days.

G) BCA protein quantification absorbance 562 nm values for uninfected and Mtb-infected (MOI=5 and 10) BMDMs at 24h post infection.

H) Oxygen consumption rate (OCR), maximal respiration, spare respiratory capacity, ATP production, and basal respiration measured by Agilent Seahorse Metabolic Analyzer in uninfected and Mtb-infected (MOI=5 and 10) BMDMs at 24h.

I) BCA protein quantification absorbance 562 nm values for uninfected and Mtb-infected (MOI=1) BMDMs at 1, 3, 5 days.

J) GSEA pathway dot plots displaying top 20 upregulated and downregulated Reactome pathways ranked by statistical significance in Mtb-infected BMDMs (MOI=1) at Day 5 versus Day 3 post-infection. Genome-wide ranking vector ordered by $\log_2$Fold Change values. Pathways are ordered on the y-axis by Normalized Enrichment Score (NES). Dot size scales proportionally with the number of core enrichment genes contributing to the pathway. Color specifies upregulation (red) or downregulation (blue). Statistical filtering was controlled using Benjamini-Hochberg adjusted p-values.

K) Hierarchical clustering heatmaps depicting MHC-II Antigen Presentation Reactome pathway gene sets filtered by custom functional effector signatures. Relative transcript abundances derived from Mtb-infected BMDMs (MOI=1) at Day 5 versus Day 3 were normalized via variance-stabilizing transformation (VST) and mapped to row-wise Z-scores (red denotes upregulation; blue denotes downregulation). Rows are organized by unsupervised hierarchical clustering, and columns are fixed by experimental sample group.

L) As in K, but depicting Interferon Signaling.

M) As in K, but Role of Phospholipids in Phagocytosis.

N) As in K, Role of Vesicle-mediated transport.

Statistical analysis: *$p < 0.05$, **$p < 0.01$, ***$p < 0.001$, ****$p < 0.0001$. Statistical differences were determined for (B-D, P) using one-way ANOVA with Tukey's post-test, (E-J, L-O) using two-way ANOVA with Sidak's post-test, and (Q) using multiple two-tailed Student's unpaired t tests.

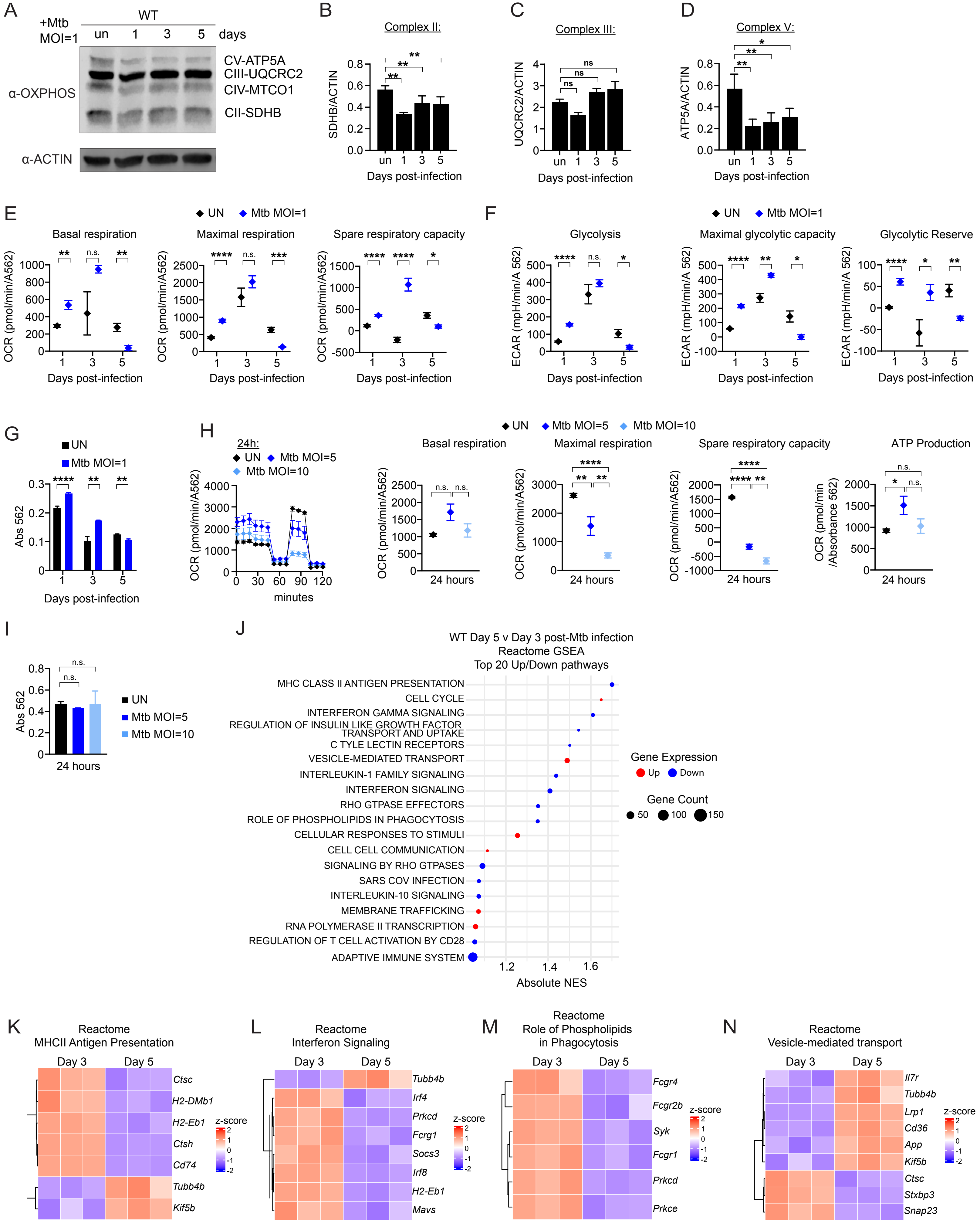

A
+Mtb MOI=1
WT
un 1 3 5 days
α-OXPHOS
CV-ATP5A
CIII-UQCRC2
CIV-MTCO1
CII-SDHB
α-ACTIN
B
Complex II:
SDHB/ACTIN
Days post-infection
C
Complex III:
UQCRC2/ACTIN
Days post-infection
D
Complex V:
ATP5A/ACTIN
Days post-infection
E
UN Mtb MOI=1
Basal respiration
Maximal respiration
Spare respiratory capacity
OCR (pmol/min/A562)
Days post-infection
F
UN Mtb MOI=1
Glycolysis
Maximal glycolytic capacity
Glycolytic Reserve
ECAR (mpH/min/A 562)
Days post-infection
G
UN
Mtb MOI=1
Abs 562
Days post-infection
H
24h:
UN Mtb MOI=5
Mtb MOI=10
OCR (pmol/min/A562)
minutes
UN Mtb MOI=5 Mtb MOI=10
Basal respiration
Maximal respiration
Spare respiratory capacity
ATP Production
24 hours
OCR (pmol/min /Absorbance 562)
I
Abs 562
UN
Mtb MOI=5
Mtb MOI=10
24 hours
J
WT Day 5 v Day 3 post-Mtb infection
Reactome GSEA
Top 20 Up/Down pathways
MHC CLASS II ANTIGEN PRESENTATION
CELL CYCLE
INTERFERON GAMMA SIGNALING
REGULATION OF INSULIN LIKE GROWTH FACTOR TRANSPORT AND UPTAKE
C TYLE LECTIN RECEPTORS
VESICLE-MEDIATED TRANSPORT
INTERLEUKIN-1 FAMILY SIGNALING
INTERFERON SIGNALING
RHO GTPASE EFFECTORS
ROLE OF PHOSPHOLIPIDS IN PHAGOCYTOSIS
CELLULAR RESPONSES TO STIMULI
CELL CELL COMMUNICATION
SIGNALING BY RHO GTPASES
SARS COV INFECTION
INTERLEUKIN-10 SIGNALING
MEMBRANE TRAFFICKING
RNA POLYMERASE II TRANSCRIPTION
REGULATION OF T CELL ACTIVATION BY CD28
ADAPTIVE IMMUNE SYSTEM
Absolute NES
Gene Expression
Up Down
Gene Count
50 100 150
K
Reactome
MHCII Antigen Presentation
Day 3 Day 5
Ctsc
H2-DMb1
H2-Eb1
Ctsh
Cd74
Tubb4b
Kif5b
z-score
L
Reactome
Interferon Signaling
Day 3 Day 5
Tubb4b
Irf4
Prkcd
Fcrg1
Socs3
Irf8
H2-Eb1
Mavs
M
Reactome
Role of Phospholipids in Phagocytosis
Day 3 Day 5
Fcgr4
Fcgr2b
Syk
Fcgr1
Prkcd
Prkce
N
Reactome
Vesicle-mediated transport
Day 3 Day 5
Il7r
Tubb4b
Lrp1
Cd36
App
Kif5b
Ctsc
Stxbp3
Snap23

**Figure S5.**

A) Basal respiration, maximal respiration, spare respiratory capacity, ATP production, and non-mitochondria respiration in untreated SCR KD and *Ndufs4* KD iBMDMs.

B) Non-glycolytic acidification, glycolysis, maximal glycolytic capacity, and glycolytic reserve in untreated SCR KD and *Ndufs4* KD iBMDMs.

C) GSEA pathway dot plots displaying the top upregulated and downregulated GO:BP pathways ranked by statistical significance in uninfected SCR KD *Ndufs4* KD versus SCR KD iBMDMs. Genome-wide ranking vector ordered by $\log_2$Fold Change values from the indicated comparison. Pathways are ordered on the y-axis by Normalized Enrichment Score (NES).

D) As in C but at 6h post-Mtb infection (MOI=5) SCR KD and *Ndufs4* KD iBMDMs.

E) As in C but at 6h post-Mtb infection (MOI=5) SCR KD and *Ndufs4* KD iBMDMs.

Statistical analysis: *p < 0.05, **p < 0.01, ***p < 0.001, ****p < 0.0001. Statistical differences were determined for (A, B) using multiple two-tailed Student’s unpaired t tests.

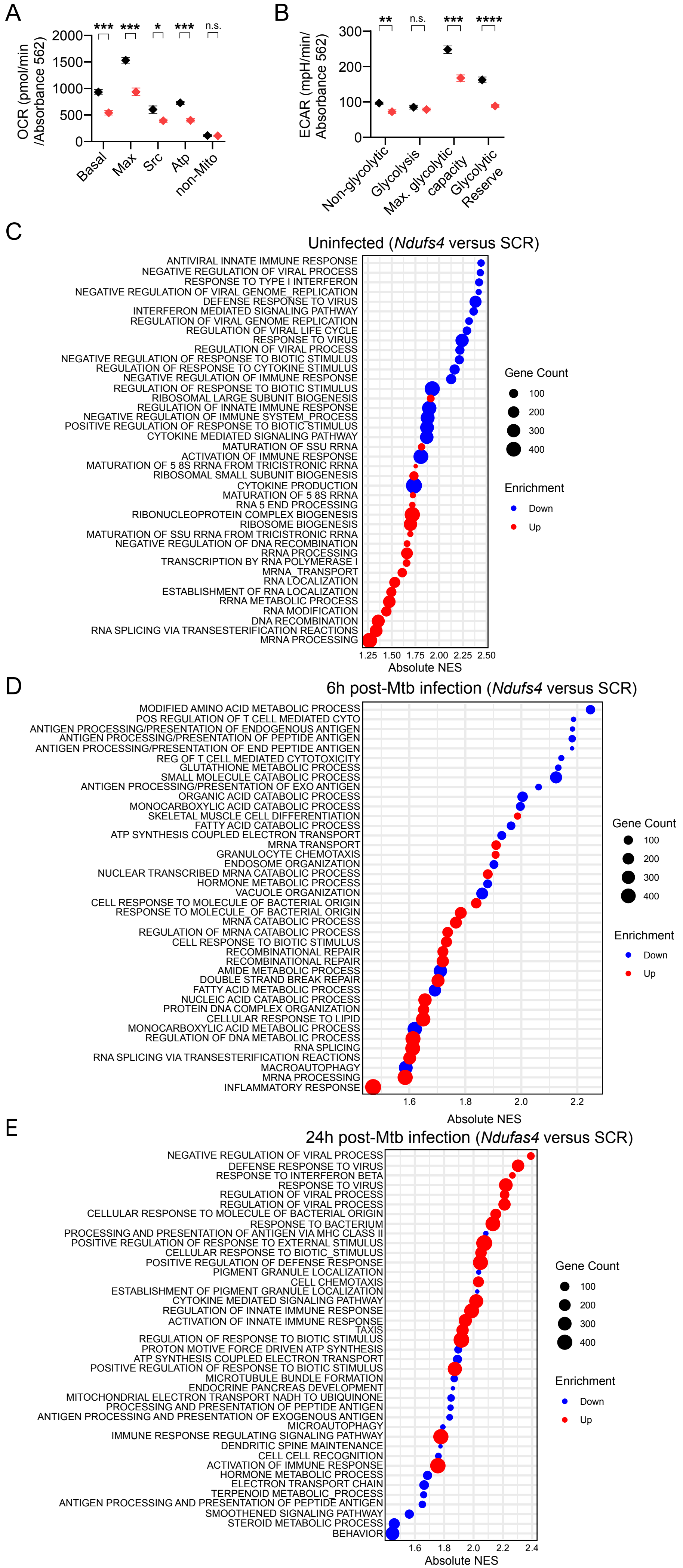

A
OCR (pmol/min /Absorbance 562)
2000
1500
1000
500
0
*** *** * *** n.s.
Basal
Max
Src
Atp
non-Mito
B
ECAR (mpH/min/ Absorbance 562)
300
200
100
0
** n.s. *** ****
Non-glycolytic
Glycolysis
Max. glycolytic capacity
Glycolytic Reserve
C
Uninfected (*Ndufs4* versus SCR)
ANTIVIRAL INNATE IMMUNE RESPONSE
NEGATIVE REGULATION OF VIRAL PROCESS
RESPONSE TO TYPE I INTERFERON
NEGATIVE REGULATION OF VIRAL GENOME_REPLICATION
DEFENSE RESPONSE TO VIRUS
INTERFERON MEDIATED SIGNALING PATHWAY
REGULATION OF VIRAL GENOME REPLICATION
REGULATION OF VIRAL LIFE CYCLE
RESPONSE TO VIRUS
REGULATION OF VIRAL PROCESS
NEGATIVE REGULATION OF RESPONSE TO BIOTIC STIMULUS
REGULATION OF RESPONSE TO CYTOKINE STIMULUS
NEGATIVE REGULATION OF IMMUNE RESPONSE
REGULATION OF RESPONSE TO BIOTIC STIMULUS
RIBOSOMAL LARGE SUBUNIT BIOGENESIS
REGULATION OF INNATE IMMUNE RESPONSE
NEGATIVE REGULATION OF IMMUNE SYSTEM_PROCESS
POSITIVE REGULATION OF RESPONSE TO BIOTIC STIMULUS
CYTOKINE MEDIATED SIGNALING PATHWAY
MATURATION OF SSU RRNA
ACTIVATION OF IMMUNE RESPONSE
MATURATION OF 5 8S RRNA FROM TRICISTRONIC RRNA
RIBOSOMAL SMALL SUBUNIT BIOGENESIS
CYTOKINE PRODUCTION
MATURATION OF 5 8S RRNA
RNA 5 END PROCESSING
RIBONUCLEOPROTEIN COMPLEX BIOGENESIS
RIBOSOME BIOGENESIS
MATURATION OF SSU RRNA FROM TRICISTRONIC RRNA
NEGATIVE REGULATION OF DNA RECOMBINATION
RRNA PROCESSING
TRANSCRIPTION BY RNA POLYMERASE I
MRNA_TRANSPORT
RNA LOCALIZATION
ESTABLISHMENT OF RNA LOCALIZATION
RRNA METABOLIC PROCESS
RNA MODIFICATION
DNA RECOMBINATION
RNA SPLICING VIA TRANSESTERIFICATION REACTIONS
MRNA PROCESSING
1.25 1.50 1.75 2.00 2.25 2.50
Absolute NES
Gene Count
100
200
300
400
Enrichment
Down
Up
D
6h post-Mtb infection (*Ndufs4* versus SCR)
MODIFIED AMINO ACID METABOLIC PROCESS
POS REGULATION OF T CELL MEDIATED CYTO
ANTIGEN PROCESSING/PRESENTATION OF ENDOGENOUS ANTIGEN
ANTIGEN PROCESSING/PRESENTATION OF PEPTIDE ANTIGEN
ANTIGEN PROCESSING/PRESENTATION OF END PEPTIDE ANTIGEN
REG OF T CELL MEDIATED CYTOTOXICITY
GLUTATHIONE METABOLIC PROCESS
SMALL MOLECULE CATABOLIC PROCESS
ANTIGEN PROCESSING/PRESENTATION OF EXO ANTIGEN
ORGANIC ACID CATABOLIC PROCESS
MONOCARBOXYLIC ACID CATABOLIC PROCESS
SKELETAL MUSCLE CELL DIFFERENTIATION
FATTY ACID CATABOLIC PROCESS
ATP SYNTHESIS COUPLED ELECTRON TRANSPORT
MRNA TRANSPORT
GRANULOCYTE CHEMOTAXIS
ENDOSOME ORGANIZATION
NUCLEAR TRANSCRIBED MRNA CATABOLIC PROCESS
HORMONE METABOLIC PROCESS
VACUOLE ORGANIZATION
CELL RESPONSE TO MOLECULE OF BACTERIAL ORIGIN
RESPONSE TO MOLECULE_OF BACTERIAL ORIGIN
MRNA CATABOLIC PROCESS
REGULATION OF MRNA CATABOLIC PROCESS
CELL RESPONSE TO BIOTIC STIMULUS
RECOMBINATIONAL REPAIR
RECOMBINATIONAL REPAIR
AMIDE METABOLIC PROCESS
DOUBLE STRAND BREAK REPAIR
FATTY ACID METABOLIC PROCESS
NUCLEIC ACID CATABOLIC PROCESS
PROTEIN DNA COMPLEX ORGANIZATION
CELLULAR RESPONSE TO LIPID
MONOCARBOXYLIC ACID METABOLIC PROCESS
REGULATION OF DNA METABOLIC PROCESS
RNA SPLICING
RNA SPLICING VIA TRANSESTERIFICATION REACTIONS
MACROAUTOPHAGY
MRNA PROCESSING
INFLAMMATORY RESPONSE
1.6 1.8 2.0 2.2
Absolute NES
Gene Count
100
200
300
400
Enrichment
Down
Up
E
24h post-Mtb infection (*Ndufas4* versus SCR)
NEGATIVE REGULATION OF VIRAL PROCESS
DEFENSE RESPONSE TO VIRUS
RESPONSE TO INTERFERON BETA
RESPONSE TO VIRUS
REGULATION OF VIRAL PROCESS
REGULATION OF VIRAL PROCESS
CELLULAR RESPONSE TO MOLECULE OF BACTERIAL ORIGIN
RESPONSE TO BACTERIUM
PROCESSING AND PRESENTATION OF ANTIGEN VIA MHC CLASS II
POSITIVE REGULATION OF RESPONSE TO EXTERNAL STIMULUS
CELLULAR RESPONSE TO BIOTIC_STIMULUS
POSITIVE REGULATION OF DEFENSE RESPONSE
PIGMENT GRANULE LOCALIZATION
CELL CHEMOTAXIS
ESTABLISHMENT OF PIGMENT GRANULE LOCALIZATION
CYTOKINE MEDIATED SIGNALING PATHWAY
REGULATION OF INNATE IMMUNE RESPONSE
ACTIVATION OF INNATE IMMUNE RESPONSE
TAXIS
REGULATION OF RESPONSE TO BIOTIC STIMULUS
PROTON MOTIVE FORCE DRIVEN ATP SYNTHESIS
ATP SYNTHESIS COUPLED ELECTRON TRANSPORT
POSITIVE REGULATION OF RESPONSE TO BIOTIC STIMULUS
MICROTUBULE BUNDLE FORMATION
ENDOCRINE PANCREAS DEVELOPMENT
MITOCHONDRIAL ELECTRON TRANSPORT NADH TO UBIQUINONE
PROCESSING AND PRESENTATION OF PEPTIDE ANTIGEN
ANTIGEN PROCESSING AND PRESENTATION OF EXOGENOUS ANTIGEN
MICROAUTOPHAGY
IMMUNE RESPONSE REGULATING SIGNALING PATHWAY
DENDRITIC SPINE MAINTENANCE
CELL CELL RECOGNITION
ACTIVATION OF IMMUNE RESPONSE
HORMONE METABOLIC PROCESS
ELECTRON TRANSPORT CHAIN
TERPENOID METABOLIC_PROCESS
ANTIGEN PROCESSING AND PRESENTATION OF PEPTIDE ANTIGEN
SMOOTHENED SIGNALING PATHWAY
STEROID METABOLIC PROCESS
BEHAVIOR
1.6 1.8 2.0 2.2 2.4
Absolute NES
Gene Count
100
200
300
400
Enrichment
Down
Up

**Figure S6**

(A) Boxplot of IGRA scores of 45 patients used in the study.

A

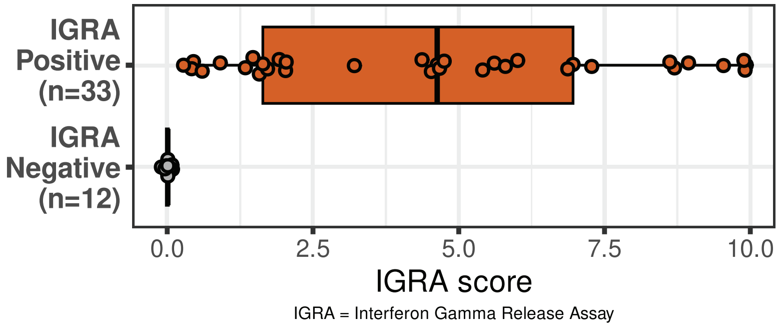
IGRA Positive (n=33)
IGRA Negative (n=12)
0.0
2.5
5.0
7.5
10.0
IGRA score
IGRA = Interferon Gamma Release Assay
